# Rapid and largely reversible shifts in the canine fecal metabolome during dietary change

**DOI:** 10.64898/2026.09.26.754668

**Authors:** Tamás Járay, Zsolt Csabai, Zsolt Boldogkői, Dóra Tombácz

## Abstract

Diet changes the fecal metabolome quickly, but it is less clear how the metabolome recovers once the original diet is restored. We followed nine Pumi dogs through an owner-managed switch from dry food to a raw (BARF) diet and back, and profiled 72 fecal samples by untargeted UPLC–MS. Diet phase accounted for roughly half of the dog-centered variation in both ionization modes. At the first sampling point after the switch to raw food, more than 13,000 LC–MS features had changed, and the return to dry food produced a response of similar size in the opposite direction: of the features that changed at both transitions, more than 99% reversed. By the last sampling point, no ESI+ features and only 13 ESI− features met the significance and effect-size thresholds for a difference from the second dry-food baseline. The overall BARF-associated pattern persisted when individual dogs were left out, and the displayed candidate associations remained significant in pedigree-adjusted models, although individual feature effects depended on normalization. Putative indole/tryptophan, bile-acid, and aromatic amino-acid metabolites showed different response and recovery patterns. In this cohort, the fecal metabolome changed rapidly during the dietary transition and returned largely toward baseline after dry food was resumed, with differences in recovery among dogs.

## Introduction

Diet determines which nutrients reach the canine gut, shapes digestive physiology and sets the conditions under which intestinal microorganisms grow. Macronutrient composition, processing, fiber content and raw feeding all influence fecal microbial composition and metabolism [1–8]. Because fecal metabolomics captures compounds of dietary, host and microbial origin, it allows these effects to be studied together.

Several studies have compared the metabolomes of raw-fed and commercially fed dogs. Schmidt et al. found distinct fecal metabolomic profiles in dogs fed BARF and commercial diets [1], Scarsella et al. used nuclear magnetic resonance spectroscopy to distinguish dogs fed extruded, homemade and raw meat-based diets [2], and Hiney et al. reported serum metabolomic differences between raw-fed and kibble-fed dogs, many of which could reflect macronutrient composition [8]. These between-dog comparisons provide limited information about how an individual dog responds to a diet change and how closely its metabolome returns to baseline afterward.

Longitudinal studies have addressed the timing of these changes. Sandri et al. showed that switching between extruded food and a vegetable-supplemented raw meat diet altered the fecal microbiota and fermentation products [3]. In 12 dogs followed after an abrupt diet change, Lin et al. saw fecal metabolite profiles shift within about 2 days, whereas the microbiota needed about 6 days to stabilize [5]. Altering the protein-to-carbohydrate ratio affects indole, amino-acid, lipid and carbohydrate metabolism [6], and a replicated 5 × 5 Latin-square study by Geary et al. found differences in fecal metabolites and microbiota among extruded, fresh, frozen raw and freeze-dried raw diets [7].

Most recently, Nearing et al. combined fecal and serum metabolomics with shotgun metagenomics and metatranscriptomics in a randomized crossover study of low-carbohydrate foods, and reported changes in tryptophan derivatives, secondary bile acids, short-chain fatty acid fermentation, amino acids, dipeptides and ammonia [9]. Together, these studies show that canine gut chemistry responds quickly to diet. Much less is known about what happens when the original diet is resumed: how fast, and how completely, the metabolome returns to its starting state.

Dogs are also a useful system for studying how gut microbes respond to diet. Comparative metagenomics has identified similarities between canine and human gut microbiomes in gene content and responses to dietary macronutrients [10], and longitudinal profiling has shown how age, diet and birth mode shape the canine microbiome over time [11]. The strengths and limits of the dog as a model for human gut microbiome research have recently been reviewed [12]. The fecal metabolomes of the two species are not interchangeable, but short-term animal-based and plant-based diets also reshape microbial composition and activity in humans [13], so response and recovery are relevant questions in both.

Here we took advantage of a diet change planned by a breeder: nine dogs moved from dry food to a raw diet and back, and we sampled their feces before, during and after the raw-feeding period. We asked how consistently the fecal metabolome responded, how closely it returned to its starting state once dry food was resumed, and how much recovery differed between dogs.

## Results

### Analytical quality and overall metabolome changes

Untargeted profiling detected 21,503 ESI+ and 24,928 ESI− features, of which 19,630 and 22,886, respectively, passed QC filtering. Median QC RSD was approximately 16.5% in ESI+ and 14.4% in ESI−, more than 90% of detected features in each polarity had QC RSD <30%, and the total-signal CV of the pooled QC injections was 2.0% and 3.0%, respectively. All longitudinal analyses used the QC-filtered matrices.

Diet phase was strongly associated with global metabolome structure. The first principal component explained 52.2% of the QC-filtered, Pareto-scaled variance in ESI+ and 44.8% in ESI−, and separated BARF samples from DRY and REDRY samples (Figure 1). With permutations restricted within dogs, phase remained strongly associated with metabolome structure (ESI+: pseudo-F=39.34, partial R²=0.563; ESI−: pseudo-F=29.42, partial R²=0.491; permutation P=0.0001 for both).

**Figure 1.**
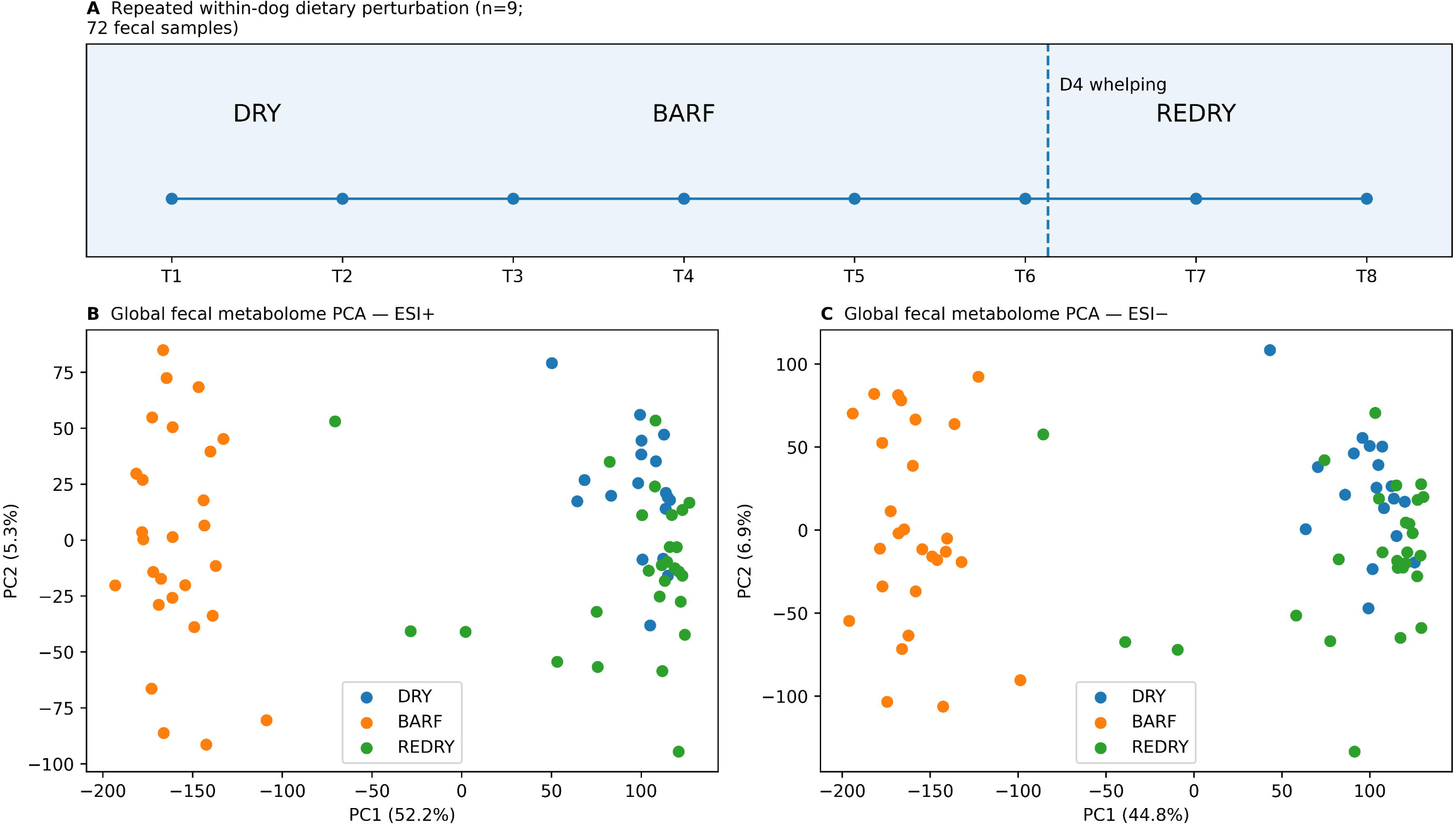
Study design and global metabolome structure. (A) Dietary sequence followed by each dog across eight sampling time points. T1–T2 were collected during the initial dry-food phase, T3–T5 during the raw-food phase, and T6–T8 after return to the original dry diet; the timing of D4 whelping is indicated. (B,C) Principal-component analysis of QC-filtered biological samples in ESI+ and ESI−, respectively, colored by diet phase. Sampling positions are schematic and not spaced proportionally to elapsed days; D4 whelping occurred two days after T6.

### Rapid changes after both diet switches

Metabolome changes were evident at the first sampling point after raw feeding began. T3, the first BARF sample, was collected two days after T2, the last dry-food sample. Between these two samples, 13,209 ESI+ and 14,045 ESI− features changed (BH-FDR <0.05 and |log2FC| ≥0.5), compared with only 2,277 and 2,023 features between the two DRY baseline samples (T1 versus T2). The first sample after the return to dry food showed a similarly large response: 13,269 ESI+ and 14,004 ESI− features changed between T5 and T6, which were also two days apart and bracketed the BARF-to-REDRY switch (Figure 2A).

**Figure 2.**
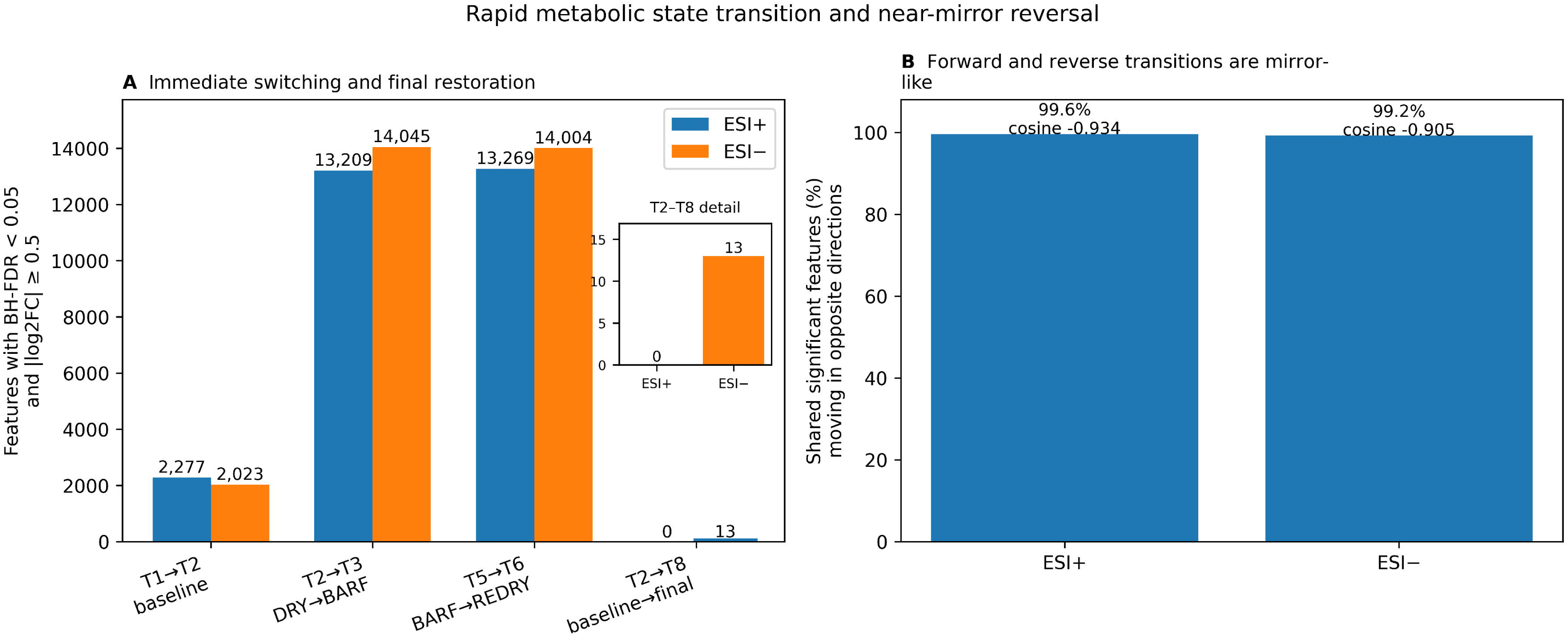
Changes after the two diet switches. (A) Numbers of QC-filtered LC–MS features meeting BH-FDR <0.05 and |log2FC| ≥0.5 for natural baseline variation (T1→T2), the forward diet switch (T2→T3), the reverse switch (T5→T6), and the baseline-to-final comparison (T2→T8). At T2→T8, 0 ESI+ features and 13 ESI− features met the threshold. The inset magnifies the T2→T8 counts without changing the heights in the main panel. (B) Percentage of features significant in both forward and reverse transitions that moved in opposite directions, together with cosine similarity of the full QC-filtered forward and reverse signed effect vectors.

### Most changes reverse after return to dry food

Feature changes after the return to dry food were largely opposite in direction to those observed after raw feeding began. Of the 11,143 ESI+ and 11,110 ESI− features significant in both the T2→T3 and T5→T6 comparisons, 99.6% and 99.2%, respectively, moved in opposite directions. Across all QC-filtered features, the cosine similarity between the cohort-level forward and reverse effect vectors was −0.934 in ESI+ and −0.905 in ESI−, where −1 would indicate a perfectly mirrored response (Figure 2B). By T8, no ESI+ feature and only 13 ESI− features differed from T2 under the same threshold. In the first 10 PCA dimensions, the displacement of each dog between T2 and T8 did not differ detectably from its observed baseline displacement between T1 and T2 (exact paired P=0.906 in ESI+ and P=0.883 in ESI−; Supplementary Figure S9), although this test was not designed to demonstrate equivalence.

### Features show different patterns of response and recovery

The phase-level screen identified 15,211 ESI+ and 16,586 ESI− robust diet-responsive features, most of which decreased during BARF. By T8, the median recovery of these features was 88.5% in ESI+ and 87.4% in ESI−, and 76.3% and 73.3%, respectively, had recovered by at least 75%. Clustering separated three temporal patterns (Figure 3): large decreases that reversed quickly, sometimes with a transient overshoot; decreases that recovered more slowly and remained displaced in early REDRY; and BARF-associated increases that mostly returned rapidly to baseline. Supplementary Figure S10 shows the 30 most variable annotated features across all samples (Supplementary Table S15), and Supplementary Figure S11 shows individual-dog trajectories of selected features, illustrating how dogs differed during REDRY (Supplementary Table S16). Both selections were made post hoc and are descriptive; the variance-based selection does not validate the chemical identities.

**Figure 3.**
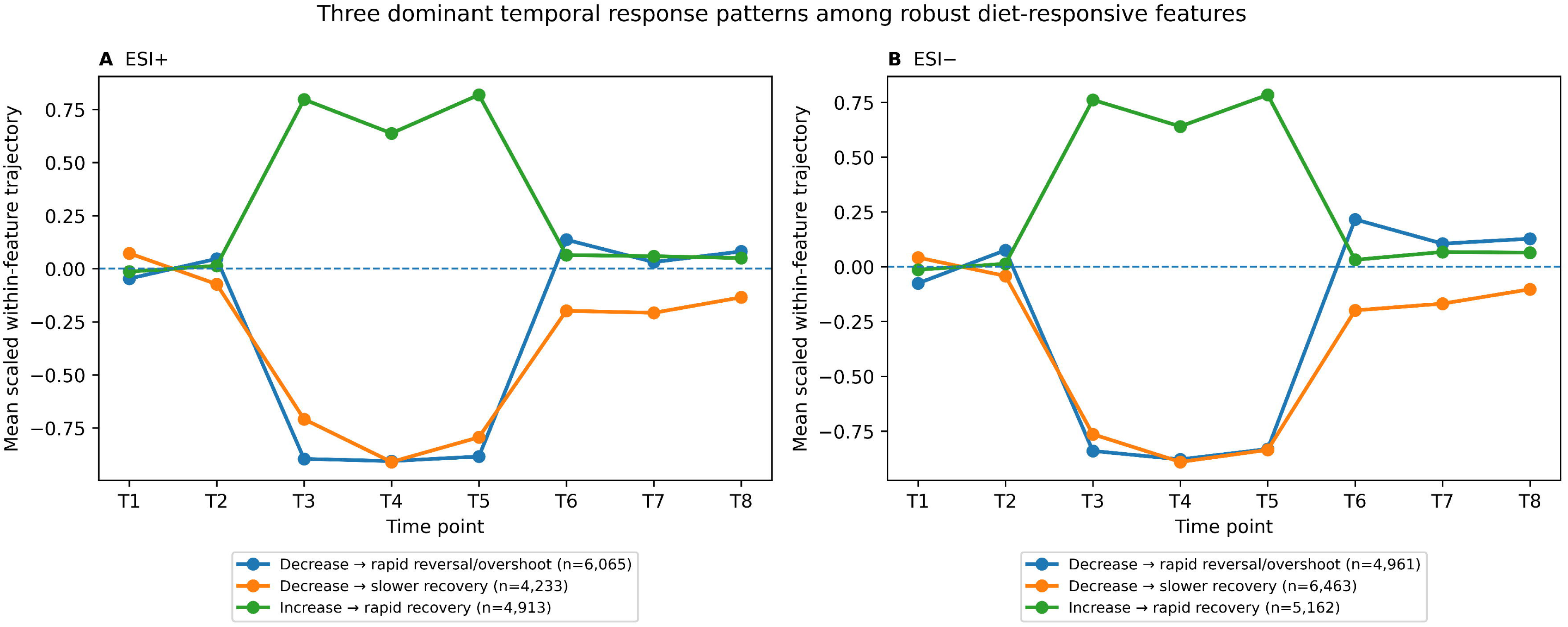
Three descriptive clusters of diet-responsive feature trajectories. (A,B) K-means clustering of baseline-centered, within-feature-scaled trajectories in ESI+ and ESI−. The three descriptive patterns comprise a BARF-associated decrease followed by rapid reversal/overshoot, a decrease followed by slower recovery, and a BARF-associated increase followed by rapid recovery. Cluster sizes are shown in the legends.

### Annotation confidence and cross-polarity support

The vendor annotation tables contained 1,510 named ESI+ and 1,553 named ESI− features, of which 1,438 and 1,488 passed QC filtering. Of these, 54 ESI+ and 43 ESI− annotations met our internal Tier A criterion, and a further 91 and 117 met Tier B. A total of 593 metabolite names occurred in both ionization modes; 147 of these matched by name, formula and retention time (within 0.05 min), and 128 of the 147 changed in the same direction during BARF. Six putative metabolites passed a stricter filter requiring a robust diet response and an mzCloud score ≥80 in both polarities: 5-hydroxyindole, genistein, 1-methyluric acid, sinapinic acid, quercetin and N-acetylsphingosine (Supplementary Figure S7; Supplementary Table S13). We treat this cross-polarity agreement as supporting evidence, not as confirmation of chemical identity.

### Patterns within selected biochemical classes

In some biochemical classes, most responsive signals moved in the same direction. In the indole/tryptophan group, 22 of 25 signals decreased during BARF in each polarity (two-sided exact binomial test with BH correction across eight classes per polarity, q=0.00125). In ESI+, all 6 bile-acid annotations increased and 19 of 26 aromatic amino-acid/phenolic annotations decreased, but neither class reached q<0.05 (q=0.0625 for each). Because LC–MS signals are correlated and identities are putative, these counts describe coordinated patterns rather than pathway enrichment or microbial origin.

### Response and recovery in the selected metabolite panel

For follow-up, we selected 13 putatively annotated features on the basis of analytical quality, annotation support, biological relevance and the initial response. Twelve of them met the BARF-versus-DRY threshold in the random-intercept mixed model after Benjamini–Hochberg correction across all 13 candidates; these are shown in Figure 4 and Table 1. The thirteenth, N-acetyltryptophan, did not meet this threshold (q=0.070) and is reported in the supplementary results. All 12 associations also held in the pedigree-adjusted model (Supplementary Table S5b). An exact sign-flip test on within-dog phase means, used as a sensitivity analysis, gave q<0.05 for all 13 candidates, including N-acetyltryptophan (q=0.035; Supplementary Table S5c), so the exact composition of the displayed set depends on the model. Because the candidates were selected and tested in the same cohort, these results are not an independent validation.

**Figure 4.**
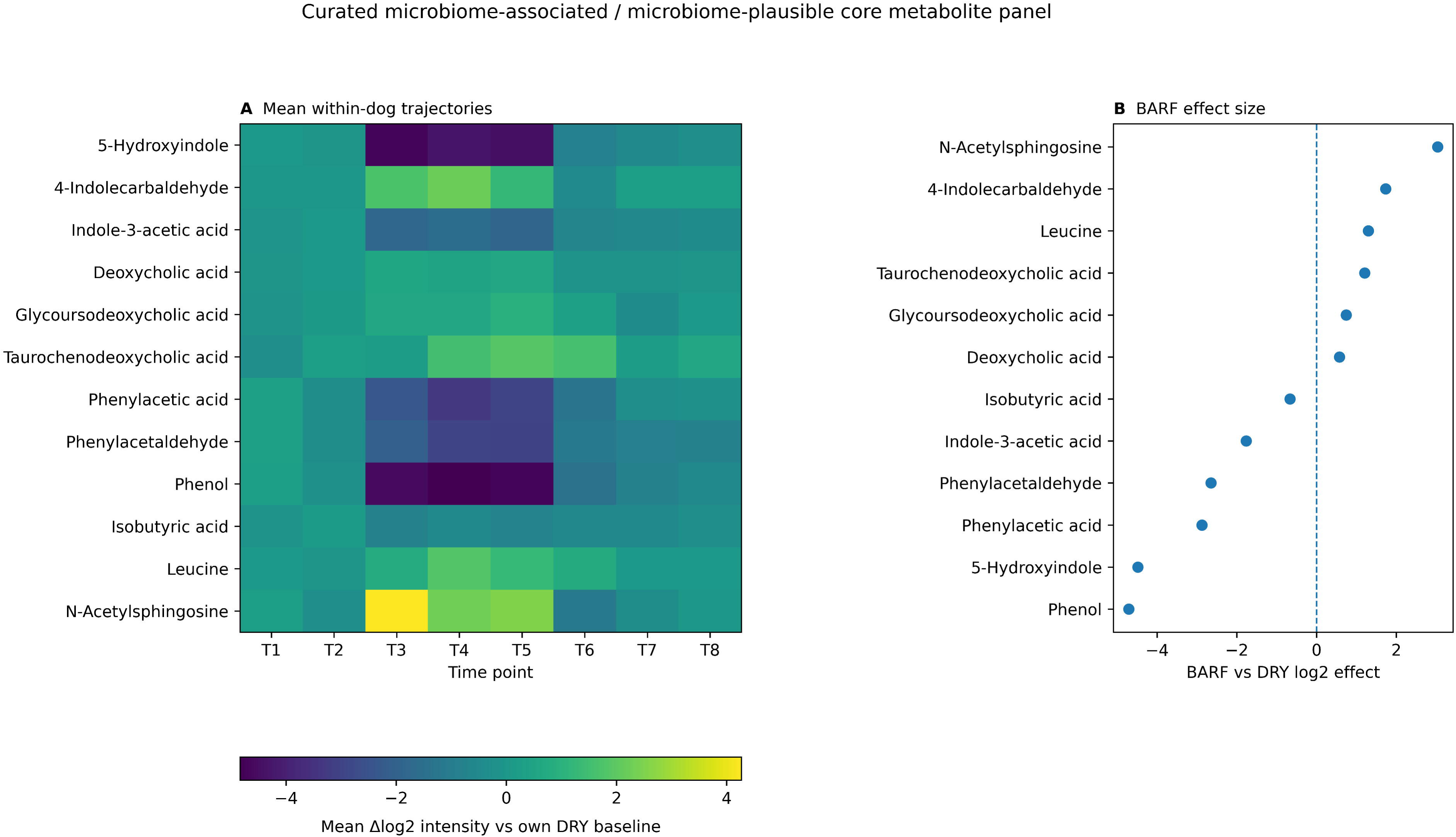
Response trajectories of selected putatively annotated features. (A) Mean within-dog trajectories of the 12 core putative metabolites relative to each dog’s dry-diet baseline across T1–T8. The horizontal color scale reports mean baseline-relative log2 intensity. (B) BARF-versus-DRY log2 effect sizes. Metabolite identities are putative; selection was based on analytical quality, annotation support, biological plausibility, and response in the initial analysis, with mixed-model results in Table 1 and exact sensitivity tests for all 13 original candidates in Supplementary Table S5c. The 12-candidate display reflects selection using the mixed-model threshold.

**Table 1.**
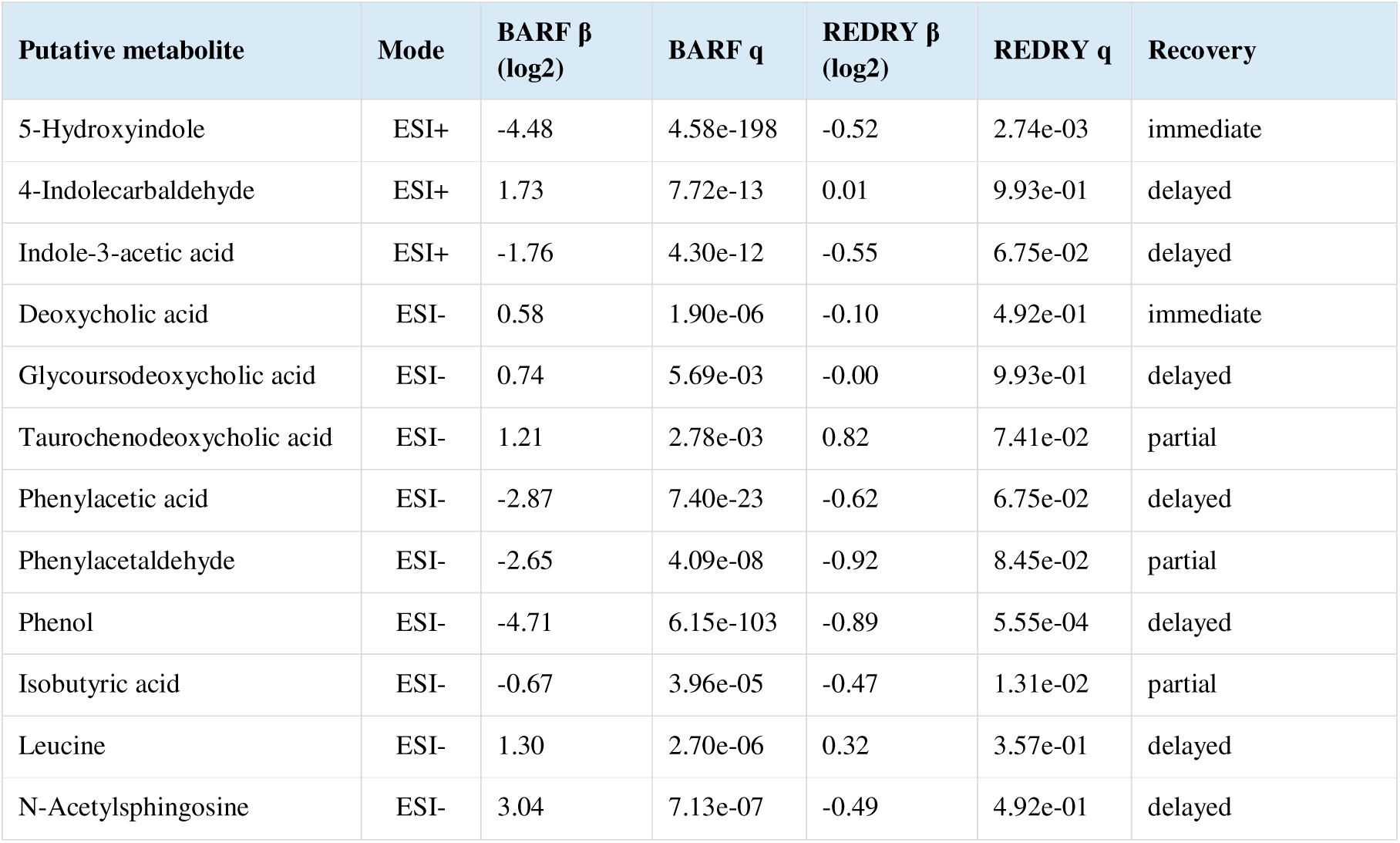
Exploratory candidate metabolite panel. Effect estimates are fixed effects from random-intercept mixed-effects models with dog as the random effect and DRY as the reference phase. q values are Benjamini–Hochberg-adjusted across the original 13 candidates separately for each contrast. The table displays the 12 candidates meeting the mixed-model BARF threshold; all 13 are retained in Supplementary Tables S5a–c. Identities are putative, and significance depends on the testing approach. Recovery labels follow the trajectory thresholds defined in Methods, not the REDRY q value.

The candidates differed in both the direction and the size of their response. Among indole-related metabolites, 5-hydroxyindole fell sharply during BARF (β=-4.48 log2) and indole-3-acetic acid also decreased (β=-1.76), whereas 4-indolecarbaldehyde increased (β=+1.73). The bile-acid candidates rose more modestly: deoxycholic acid (β=+0.58), glycoursodeoxycholic acid (β=+0.74) and taurochenodeoxycholic acid (β=+1.21). Phenylacetic acid, phenylacetaldehyde and phenol decreased, whereas leucine and N-acetylsphingosine increased. During REDRY, most candidates moved substantially back toward baseline, but not all of them completely. Three of the 12 still differed between REDRY and DRY in the mixed model after FDR correction, and four in the exact test, including taurochenodeoxycholic acid (q=0.020); N-acetyltryptophan also met the REDRY threshold in the exact test (q=0.013), giving 5 of the original 13 candidates. The number of residual differences therefore depends on the test used.

### Recovery differs among dogs

At the first REDRY sample, most dogs had already moved back toward their DRY position along the main BARF-response axis, and directional recovery approached 1.0 by T8 in most of them (Figure 5A,B). Residual distance from the DRY centroid indicated less complete recovery: several dogs remained displaced in other dimensions (Figure 5C,D), so return along the main response axis did not mean that the whole multivariate profile had recovered. D4 had the least complete recovery at T8 in both ionization modes. Its REDRY phase coincided with late pregnancy, whelping and the early postpartum period, which we examined in separate sensitivity analyses.

**Figure 5.**
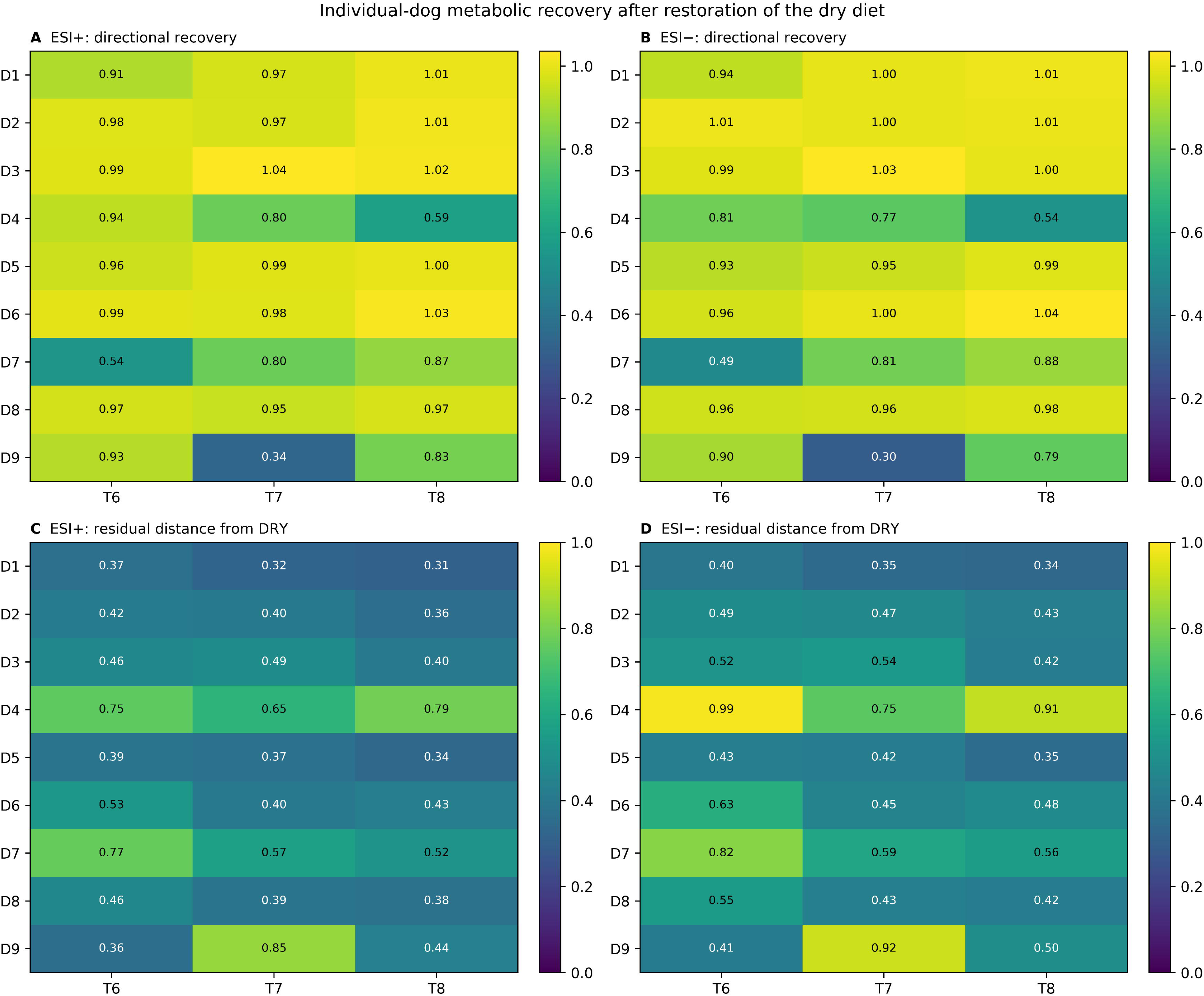
Individual-dog metabolic recovery after restoration of the dry diet. (A,B) Directional recovery for D1–D9 across T6–T8 in ESI+ and ESI−, respectively. Values near 1 indicate return toward the individual dry-diet position along the dominant BARF-response axis. (C,D) Residual multivariate distance from the individual dry-diet centroid, normalized to the original DRY→BARF displacement, in ESI+ and ESI−. D4 whelped two days after T6; its REDRY trajectory therefore overlaps the peripartum/postpartum transition. The same color limits are used across polarities for each recovery metric.

### Sensitivity to dog exclusion and analytical choices

The overall BARF-associated pattern persisted in each leave-one-dog-out analysis. Excluding D4 preserved the overall pattern, and all 12 displayed candidates kept their BARF association in the D4-excluded mixed model. In leave-one-dog-out analyses with the same exact sign-flip criteria, BARF-effect correlations were at least 0.9982 and robust-feature Jaccard overlap was at least 0.924. Removing flagged samples and recomputing within-dog phase means gave correlations of 0.99956– 1.00000 (removing REDRY-only samples cannot change the BARF-versus-DRY contrast). Normalization mattered more. PQN preserved the ranking of features but shifted BARF effects by approximately +1.00 log2 in ESI+ and +0.96 log2 in ESI−; direction agreement with vendor normalization was 88.0% and 85.6%, and robust-feature Jaccard overlap was 0.708 and 0.676. Both normalizations showed widespread diet-associated change, but the magnitude, direction and membership of individual features were normalization-dependent (Supplementary Figure S4; Supplementary Tables S6a–d).

### Exploratory peripartum trajectories in D4

Because D4’s REDRY phase overlapped late pregnancy, parturition and the early postpartum period, we examined its trajectories separately. At T6–T8, several core metabolite signals in D4 departed markedly from the median trajectory of the other eight dogs, including phenol, 5-hydroxyindole, indole-3-acetic acid, isobutyric acid, phenylacetic acid and bile-acid candidates. With a single animal, these differences cannot be attributed to reproductive status; we therefore assessed the BARF associations without D4 and interpret its recovery separately.

## Discussion

### Response and recovery after dietary change

Following the same dogs through both diet switches allowed us to measure not only how the fecal metabolome responded to raw feeding but also how it recovered. The response was fast: at the first BARF sampling point, collected two days after the last dry-food sample, more than 13,000 features in each polarity differed from the preceding dry-food sample, six to seven times the number that differed between the two baseline samples under the same threshold. The return to dry food produced a response of similar size, and more than 99% of the features that changed at both transitions moved in opposite directions. By the end of follow-up, no ESI+ and only 13 ESI− features met the significance and effect-size thresholds for a difference from T2. Final displacement did not differ detectably from the observed T1-to-T2 baseline displacement. These findings indicate substantial movement back toward baseline, but do not establish statistical equivalence or complete recovery. The opposite directions of the two responses are less consistent with a simple monotonic time trend. They do not, however, rule out other time-related changes or technical effects aligned with diet phase.

These numbers refer to LC–MS features, not distinct metabolites. Untargeted analyses generate tens of thousands of signals, including isotope peaks, adducts, fragments of the same compound and non-biological signals; in one *Escherichia coli* dataset, detailed annotation reduced approximately 25,000 features to fewer than 1,000 unique metabolites [14]. The corresponding ratio in canine feces is unknown, but the feature counts clearly do not represent thousands of independent metabolic changes.

### Comparison with previous canine studies

Our results agree with earlier comparisons of raw-fed and commercially fed dogs, which found distinct fecal [1,2] and serum [8] metabolomic profiles, the latter partly attributed to macronutrient composition. By following the same animals, we reduced confounding by stable differences between dogs and could also examine what happened once the original diet was resumed.

The timing of the observed response is consistent with the findings of Lin et al., who saw fecal metabolite profiles change within about 2 days of an abrupt diet transition [5]. Their study followed adaptation to a fiber-enriched or protein-rich diet; we additionally sampled after the return to the original food, which allowed us to quantify the reversal. This recovery phase is the main new contribution of the present study.

### Comparison with the low carbohydrate multiomics study

Nearing et al. studied 35 dogs in a randomized crossover of low-carbohydrate foods in which fat or protein replaced carbohydrate energy, combining fecal and serum metabolomics with fecal shotgun metagenomics and metatranscriptomics [9]. They reported changes in tryptophan derivatives, secondary bile acids and short-chain fatty acid fermentation, together with lower fecal amino acids and dipeptides and higher ammonia. Our putative annotations include members of several of these classes, such as indole-3-acetic acid, deoxycholic acid, isobutyric acid and amino acids. Differences in diets, analytical platforms and annotation confidence make a direct comparison difficult, however, and a shared name does not guarantee that the same compound was measured. The signal annotated as leucine, for example, increased in our data, while other amino-acid annotations behaved differently, and we did not measure ammonia.

### Possible sources of the measured compounds

Several responsive putative metabolites belong to classes linked to gut microbial activity, including indole/tryptophan derivatives, bile acids, aromatic amino-acid metabolites and fermentation products. Indole/tryptophan signals decreased predominantly in both polarities, whereas the ESI+ bile-acid and aromatic/phenolic patterns did not survive multiple-testing correction. Class membership alone cannot establish microbial origin. The raw ration contained animal protein, plant components and supplements, and any of the measured compounds could derive from the food, from host metabolism, from microbial activity or from a combination of these. We did not analyze the foods or supplements by LC–MS.

This uncertainty applies particularly to lipid- and marine-associated signals, because the raw ration contained salmon and a marine-derived supplement; these signals cannot be assigned specifically to the host or the microbiota. Paired metagenomic and metatranscriptomic data from the same samples could help test whether the metabolite trajectories track particular microbial genomes, genes or transcriptional activity.

BARF is not equivalent to any specific human diet. Raw feeding changes processing, ingredients and supplementation at the same time, so it is not a validated model of carnivore, ketogenic or Paleolithic diets in humans. What may be comparable across species is how gut chemistry responds to a change in dietary substrates and then recovers. David et al. observed rapid changes in microbial composition and activity during an animal-based diet in humans [13], and some shifts in substrate availability, fermentation and bile-associated processes may be broadly similar in dogs. Our data do not address human health effects, safety or personalized dietary recommendations.

### Individual recovery and relatedness

Leaving out one dog at a time barely changed the initial BARF response, but recovery varied among dogs. D4 was pregnant from T1 to T6 and whelped during REDRY; excluding it preserved the main BARF patterns, but within D4 the dietary and reproductive effects cannot be separated. The pedigree-adjusted random-intercept models retained BARF q<0.05 for all 12 displayed candidates, and the BARF and REDRY coefficients changed little, as expected for within-dog contrasts in a balanced, complete dataset. This adjustment does not make the dogs independent or account for correlated individual responses to diet, and several variance estimates reached their boundaries. With nine related dogs, we cannot reliably estimate heritability or predict how dogs in the wider population would respond.

### Limitations

The study has several limitations. The nine dogs were related, belonged to one breed and lived on one farmstead, which limits generalization. All dogs followed the same owner-managed sequence, without randomization, counterbalancing or a concurrent control group kept on the original diet, so diet phase coincided with calendar time. The raw diet was fed together with supplements, and the results therefore concern the complete feeding regimen. Sex and housing were confounded, one dog was pregnant and gave birth during follow-up, food intake was only approximate, and the foods themselves were not analyzed.

Some details were not available from the analytical provider, including parts of the preprocessing workflow, the injection order, pooled-QC preparation and annotation tolerances. Without injection-order and batch information, we could not fully exclude technical effects aligned with diet phase; tight clustering of the QC samples alone does not rule this out. Adducts and isotopes could not be fully resolved, and metabolite identities remain putative. The measured intensities are relative signals, not absolute concentrations, and normalization changed the magnitude and, for some features, the direction of individual effects. Candidate metabolites were selected and tested in the same dogs, candidate-level significance differed between the mixed model and the exact test, and the class-level binomial tests did not account for dependence among LC–MS signals. Finally, failure to detect a residual difference is not evidence of equivalence, and recovery estimates describe the sampled time points rather than a stable trait of individual dogs.

The pedigree was compiled from breeder-supplied records of limited depth and still requires registry-level verification; undocumented relationships among founders could not be recovered. The pedigree models accounted for covariance between dog intercepts, but not for relatedness-dependent responses to diet or for confounding by calendar time. Likewise, permutations blocked within dogs preserve the repeated-measures structure but do not establish temporal exchangeability or account for relatedness, and exact sign-flip tests assume symmetric, independent dog-level contrasts. These assumptions should be kept in mind when interpreting P values and FDR estimates in a small, nonrandomized cohort.

### Conclusions

The fecal metabolome differed from the preceding dry-food sample at the first sampling point during raw feeding and moved largely back toward baseline after the original diet was resumed. Recovery was substantial by the end of follow-up but varied among dogs. Sampling the same animals through both transitions allowed us to describe the initial response and subsequent recovery. Larger, controlled studies are needed to determine how reproducible these individual recovery patterns are. The present data do not support personalized feeding recommendations.

## Methods

### Study design, animals, and sampling

Nine Pumi dogs were followed during a dietary change independently planned and managed by their breeder/owner: an initial dry-food phase (DRY), an approximately one-month raw-food period (BARF), and a return to the original dry diet (REDRY). The dogs lived in a shared rural farmstead environment in the Southern Great Plain region of Hungary with two owners. The breeder decided the feeding regimen, quantities, and start date; investigators scheduled noninvasive fecal collections around these owner decisions and did not initiate or prescribe the dietary change. Eight sampling time points comprised T1–T2 during DRY, T3–T5 during BARF, and T6–T8 during REDRY, yielding 72 samples from nine dogs. The dog was the biological unit for within-dog contrasts; repeated samples were not treated as 72 independent animals. Dogs were designated D1–D9 in descending age order, with sex, housing, pedigree, and reproductive status recorded separately (Table 2). The cohort comprised four males and five females. Seven adult dogs were aged 1.75–8.05 years at T1, alongside one younger dog aged 0.81 years and one older dog aged 10.66 years. The single-breed, related cohort and shared husbandry setting reduced variation in background characteristics relative to a heterogeneous multi-household cohort. Age, sex and housing were retained as descriptive metadata rather than analyzed as separate explanatory factors; housing details are reported in Table 2.

**Table 2.** Anonymized study-dog metadata. Study identifiers D1–D9 are ordered from oldest to youngest at study initiation.

| Study ID | Sex | Age at T1 (y) | Housing |
| --- | --- | --- | --- |
| D1 | M | 10.7 | outdoor continuously |
| D2 | F | 8.0 | outdoor daytime / indoors at night |
| D3 | M | 7.7 | outdoor continuously |
| D4 | F | 6.6 | outdoor daytime / indoors at night |
| D5 | F | 6.1 | outdoor daytime / indoors at night |
| D6 | M | 5.5 | outdoor continuously |
| D7 | F | 5.0 | outdoor daytime / indoors at night |
| D8 | F | 1.7 | outdoor daytime / indoors at night |
| D9 | M | 0.8 | outdoor continuously |

D4 was pregnant during T1-T6 and whelped on March 23, 2024. Accordingly, T5 was 4 days prepartum, T6 was 2 days prepartum, T7 was 13 days postpartum, and T8 was 27 days postpartum. D6 was the sire of the litter. D4 was not isolated from the other dogs during the reproductive transition. Sensitivity analyses excluding D4 assessed BARF-associated feature effects, candidate-metabolite associations and the global phase association.

Ethics statement. Only spontaneously voided fecal samples were collected; no invasive sampling or investigator-directed intervention was performed. The breeder independently determined the dietary change, its timing, diet composition, and feeding amounts. Investigators aligned sample collection with those decisions and did not direct feeding, housing, or veterinary care. Following written consultation with Dr. József Kaszaki regarding this noninterventional, noninvasive sampling protocol, the authors were advised that formal animal-experiment approval was not required. The owner consented to participation in the study and collection of fecal samples.

Fecal samples were frozen in liquid nitrogen and subsequently stored at −80 °C until use.

### Dietary exposure and husbandry

Dogs were fed once daily in the evening throughout the study. All decisions concerning feed selection, ration composition, and the amount offered to each dog were made exclusively by the breeder/owner; the research team did not provide dietary products or prescribe feeding amounts. During DRY and REDRY, dogs received a commercial tuna-and-rice dry food at approximate daily amounts of 250 g for males and 150 g for females. During BARF, dogs received a commercial frozen raw ration at approximately 225-250 g/day for males and approximately 200 g/day for females. Each dog also received approximately one tablespoon of a dried vegetable supplement and a small spoonful of green-lipped mussel powder during the BARF phase.

The dry food contained 25% crude protein and 13% crude fat (3,910 kcal/kg). The raw ration contained 14% crude protein, 20.8% crude fat, 4.6% crude ash, 0.8% crude fiber, and 59.8% moisture (244 kcal/100 g).

Housing remained unchanged across diet phases. Males lived outdoors continuously; females were outdoors during the day and indoors overnight. Sex and housing were therefore confounded. We retained these variables as descriptive metadata and did not test them as separate explanatory factors. The dogs shared a breed and farmstead environment, reducing differences in their background conditions.

### Pedigree structure

Pedigree records for the nine dogs, supplied by the breeder, were merged into a pedigree of 238 animals, including 142 animals with documented parents and 96 assumed founders. Repeated ancestors were represented once, with parentage linked across screenshots; the registered name Cuidado Xotic was used for D2. Unknown-parent founders were assumed unrelated and noninbred, so all coefficients are conditional on the documented pedigree depth. The numerator relationship matrix A was calculated in parent-before-offspring order, with Aij = (Asire,j + Adam,j)/2 for off-diagonal entries and Aii = 1 + Asire,dam/2. Thus, Aii = 1 + Fi and the kinship matrix is A/2. All 36 distinct pairwise comparisons among the nine study dogs (9 × 8 / 2) had a positive documented additive relationship. D6 and D7 were first cousins through their full-sibling parents, with additional shared ancestry (A = 0.137772); D4 and D6 also shared ancestors (A = 0.004395). Their mating was recorded separately as reproductive metadata. Study-dog F estimates ranged from 0 to 0.044189, conditional on the founder assumptions. The parentage table and the study-dog matrix are supplied in Supplementary Tables S14a and S14b, with identifier reconciliation in Supplementary Table S14c and an overview in Supplementary Figure S1.

### Untargeted fecal metabolomics

Untargeted metabolomics was performed by Creative Proteomics on 72 fecal samples using a Vanquish Flex UPLC system coupled to a Q Exactive Plus high-resolution mass spectrometer. Samples were extracted with methanol at 8 µL/mg raw material, homogenized with two 5 mm metal balls for 180 s at 65 Hz twice, sonicated for 30 min at 4°C, incubated for 1 h at −20°C, vortexed for 30 s, and centrifuged for 10 min at 12,000 rpm and 4°C. A 200-µL aliquot of supernatant was supplemented with DL-o-chlorophenylalanine internal standard and filtered through a 0.22-µm membrane before LC-MS analysis.

Chromatographic separation used an ACQUITY UPLC HSS T3 column (100×2.1 mm, 1.8 μm). Mobile phases were 0.05% formic acid in water (A) and acetonitrile (B), using a 16-min gradient at 0.3 mL/min and 40°C. Data were acquired in both positive and negative heated-electrospray-ionization modes over m/z 70–1050. Ten pooled/technical QC injections (QC1–QC10) were included in each polarity according to the vendor-supplied data matrices.

### Preprocessing, analytical QC, and normalization sensitivity

Vendor-supplied raw-intensity, normalized-intensity, and annotation tables were analyzed separately by ionization mode. For global feature analyses, features with RSD <30% calculated from raw intensities across 10 QC injections were retained (sample standard deviation, ddof=1). The annotated-feature master table used the corresponding RSD calculated from vendor-normalized QC intensities. These filters are not identical: four annotated rows retained in Supplementary Table S2 fail the raw-QC threshold. The additional exploratory heatmaps use the intersection of the annotated table and the raw-QC-passing feature set. Vendor normalization was reconstructed as each raw feature intensity divided by the total intensity of all features in that sample and multiplied by 1,000,000 (maximum absolute discrepancy <3 × 10−9). For PQN, the reference was the featurewise median raw intensity across all 72 samples among QC-passing features; each sample was divided by the median of its featurewise quotients to that reference. Raw and normalized BARF-effect vectors were also compared; their Pearson and Spearman correlations were approximately 1, indicating concordance of relative feature effects rather than proving equality of absolute effects. Pooled-QC total-signal CV was 2.0% in ESI+ and 3.0% in ESI−. For Supplementary Figure S2, each QC injection was total-signal normalized over the original full feature matrix before QC-passing features were selected, matching the normalization universe used for biological samples. For Supplementary Figure S10, exactly 30 unique mode–feature identifiers were selected by descending log2-intensity variance across 72 samples, with feature identifier as a deterministic tie-breaker. Feature-wise z-scores used the sample standard deviation across all 72 log2 intensities. The display range was fixed at −3 to +3; values outside this range were retained in the source data. No signals were merged solely because they shared a putative name. Supplementary Figure S11 plots the supplied normalized intensities for ESI+ features 746, 12299 and 1501 separately for each dog, selected post hoc to illustrate response direction and individual variability. No additional imputation or zero replacement was applied; chemical names remain putative. Sample-wise multiplicative normalization adds a common shift to mean log2 contrasts across features. A perfect correlation between raw and normalized effect vectors therefore does not independently establish normalization robustness.

### Global multivariate and feature-level longitudinal analysis

QC-filtered intensities were log2-transformed and Pareto-scaled for global principal-component analysis (PCA). Association between diet phase and multivariate metabolome structure was tested separately in ESI+ and ESI− using Euclidean distances in log2-transformed, Pareto-scaled space, with dog effects removed and 10,000 permutations of phase labels within each dog (seed 20260923). The pseudo-F statistic used 2 phase and 61 residual degrees of freedom, partial R² was the fraction of dog-centered sum of squares explained by phase, and Monte Carlo p values used (exceedances + 1)/(10,000 + 1). These blocked permutations retain the dog structure but do not independently adjust for serial correlation or pedigree; phase effects were summarized by pseudo-F statistics and partial R². At the feature level, the primary BARF-versus-DRY effect was the mean within-dog difference between the three BARF samples (T3–T5) and the two DRY samples (T1–T2). Significance was assessed with an exact two-sided sign-flip permutation test across the nine dogs (2^9=512 sign configurations), followed by Benjamini–Hochberg false-discovery-rate correction within each ionization mode. A feature was classified as a robust diet-responsive signal only when q<0.05, |BARF–DRY log2 effect|≥0.5, and at least 7 of 9 dogs changed in the same direction. In the annotated-metabolite layer, BH adjustment was performed separately among the 1,438 ESI+ and 1,488 ESI− QC-passing named annotations; these q values and robust flags were used for the biochemical-class summaries and differ from the full-feature screening family. No nonpositive normalized biological intensities occurred in the QC-passing matrices.

### Adjacent-transition, reversal, and temporal-cluster analyses

To examine changes after each diet switch, paired within-dog feature-level comparisons were performed for T1 versus T2 (natural baseline variability), T2 versus T3 (forward DRY-to-BARF transition), T5 versus T6 (reverse transition), and T2 versus T8 (baseline-to-final comparison). For each comparison, log2 intensity differences across dogs were tested with paired t tests, with Benjamini–Hochberg correction applied across features separately for each comparison and ionization mode; transition counts used q<0.05 together with |log2FC|≥0.5. Forward/reverse symmetry was quantified from the complete QC-filtered feature vectors using cosine similarity between T2→T3 and T5→T6 effects, and by the fraction of features significant in both transitions that changed in opposite directions. To describe patterns over time, robust diet-responsive features were centered to the individual dry-diet baseline, scaled within feature, and clustered by k-means (k=3; random_state=42; n_init=50). Final-state displacement was compared with baseline variability using Euclidean distances in the first 10 PCA dimensions, calculated with the deterministic full-SVD solver for T1→T2 and T2→T8; the paired dog-level distance difference was evaluated with an exact two-sided sign-flip test over all 512 sign configurations.

### Recovery kinetics

For each robust diet-responsive feature, a residual ratio was calculated at T6, T7, and T8 as the cohort mean of within-dog baseline-centered log2 displacements at that time point divided by the cohort mean BARF-versus-DRY log2 displacement; this was a ratio of cohort means, not a mean of individual ratios. A residual ratio of 1 therefore indicated persistence of the BARF-state displacement, 0 indicated return to baseline, and negative values indicated reversal beyond baseline. Recovery classes were defined from these ratios: immediate recovery, |rT6|≤0.25 and |rT8|≤0.25; delayed recovery, |rT6|>0.25 and |rT8|≤0.25; partial recovery, 0.25<rT8<0.75; persistent response, rT8≥0.75; and overshoot, rT8<−0.25. A continuous recovery fraction was calculated as 1−|rT8| and was used descriptively at the feature level. At the dog level, calculations used the full QC-passing, log2-transformed, Pareto-scaled feature space, with scaling estimated across all 72 samples. Directional recovery was quantified by projecting each REDRY metabolome onto that dog’s DRY-to-BARF response axis, with values near 1 indicating return toward the DRY position along the dominant response direction. A complementary Euclidean residual-distance metric quantified multidimensional displacement from the individual DRY centroid relative to the original DRY-to-BARF displacement.

### Metabolite annotation confidence and biochemical panels

Vendor metabolite names were treated as putative annotations rather than confirmed identities. In keeping with Metabolomics Standards Initiative principles [15], no feature was classified as identification level 1 because authentic standards and independent retention-time confirmation were not available. We therefore used an internal confidence scheme for prioritization only: Tier A, QC RSD <30% and library-match score >=90; Tier B, QC RSD <30% and score 80-89.9; and Tier C, a named QC-passing annotation without a score >=80. A ’+’ suffix was added when the same putative metabolite was observed in both ionization modes with the same name and formula, retention-time difference <=0.05 min, and concordant BARF-effect direction.

Curated biochemical classes comprised indole/tryptophan, bile-acid, aromatic amino-acid/phenolic, amino-acid, SCFA/BCFA/fermentation, histidine/bioactive-amine, sphingolipid/carnitine/choline, and purine/pyrimidine annotations. Among robust responsive annotations in each class, we used a two-sided exact binomial test against equal probabilities of increase and decrease. Benjamini–Hochberg correction was applied across eight classes separately per ionization mode. The reported dominant direction was descriptive, not a direction selected for a one-sided test. Correlated or redundant LC–MS signals may violate the binomial independence assumption, so these tests were treated as exploratory feature-level summaries rather than pathway enrichment or independently confirmed metabolite-level evidence.

### Follow-up modeling and sensitivity analyses

For each candidate metabolite, log2-normalized intensity was modeled as log2(intensity) ∼ diet phase + (1|dog), with DRY as the reference phase and dog as a random intercept. Fixed-effect coefficients therefore represent BARF-versus-DRY and REDRY-versus-DRY log2 differences. Benjamini– Hochberg correction was applied across the 13-member candidate panel separately for the two contrasts; N-acetyltryptophan did not meet this model’s BARF threshold and was omitted from the 12-candidate display, but retained in the complete supplementary results. Pedigree sensitivity used V = sigma_a^2 ZAZ′ + sigma_d^2 ZZ′ + sigma_e^2 I, where Z maps samples to dogs and A is the expanded numerator relationship matrix. All three nonnegative variance components were refitted by maximum likelihood. Residual scale was profiled analytically while the two variance ratios were optimized with L-BFGS-B using seven joint starts plus two explicit one-component boundary fits. Fixed effects were estimated by GLS, with plug-in normal-approximation Wald tests and 95% intervals; variance-component uncertainty is not included in these intervals. Optimization and likelihood checks are reported in Supplementary Table S5b. This model adjusts covariance of dog intercepts, not covariance of dog-specific phase effects. Additional sensitivity analyses excluded D4, repeated the analysis after independent probabilistic quotient normalization, removed phase-specific PCA outlier samples flagged by a robust within-phase distance z score >3.5, and sequentially omitted each dog. Leave-one-dog-out stability was summarized by Pearson correlation of BARF-effect vectors and Jaccard overlap under the same exact sign-flip screen as the primary analysis; eight-dog screens enumerate 256 sign configurations and require concordant direction in at least seven dogs. All multiple-testing adjustments were performed within the selected family of tests described above. The random-intercept models were fitted by maximum likelihood, with asymptotic two-sided Wald tests based on the normal distribution. Optimizers and convergence status are reported in Supplementary Table S5a; reproduced standard errors and 95% Wald confidence intervals are also provided. With nine dogs, these asymptotic intervals and tests require caution. As a sensitivity analysis, we also tested the 13 original candidates using within-dog phase means. For each candidate, the mean of T3–T5 or T6–T8 was compared with the mean of T1–T2 on the log2 scale. Two-sided exact sign-flip tests enumerated all 512 dog-level sign configurations. BH correction was applied across the 13 candidates separately for BARF and REDRY (Supplementary Table S5c).

### Use of artificial intelligence tools

ChatGPT (OpenAI) and Claude (Anthropic) were used to assist with language editing, code development and checking, and critical review of the manuscript, including suggestions on analytical approaches and interpretation. All suggestions were evaluated and verified by the authors and, where adopted, implemented by them. The authors take full responsibility for the content of the manuscript.

## Acknowledgments

We thank Gabriella Kassai, breeder of Serteperti Pumi Kennel (https://serteperti.hu/), for her cooperation and support with the study dogs. We thank Gábor Gulyás and Ákos Dörmő for assistance with fecal sample collection.

## Author contributions

Z.C. contributed to investigation, fecal sample collection, and coordination of the metabolomics facility. T.J. contributed to investigation, fecal sample collection, and data curation. Z.B. contributed to scientific interpretation and writing - review and editing. D.T. conceived and supervised the study, led methodology, formal analysis, data interpretation, visualization, project administration, and writing - original draft, and contributed to writing - review and editing. All authors reviewed and approved the final manuscript.

## Additional information

Competing interests. The authors declare no competing interests.

## References

1. Schmidt, M. et al. The fecal microbiome and metabolome differs between dogs fed Bones and Raw Food (BARF) diets and dogs fed commercial diets. PLoS ONE 13, e0201279 (2018). 10.1371/journal.pone.0201279

2. Scarsella, E., Segato, J., Zuccaccia, D., Swanson, K.S. & Stefanon, B. An application of nuclear magnetic resonance spectroscopy to study faecal canine metabolome. *Ital*. J. Anim. Sci. 20, 887–895 (2021). 10.1080/1828051X.2021.1925602

3. Sandri, M., Dal Monego, S., Conte, G., Sgorlon, S. & Stefanon, B. Raw meat based diet influences faecal microbiome and end products of fermentation in healthy dogs. BMC Vet. Res. 13, 65 (2017). 10.1186/s12917-017-0981-z

4. Martínez-López, L.M. et al. Effect of sequentially fed high protein, hydrolyzed protein, and high fiber diets on the fecal microbiota of healthy dogs: a cross-over study. *Anim*. Microbiome 3, 42 (2021). 10.1186/s42523-021-00101-8

5. Lin, C.-Y. et al. Longitudinal fecal microbiome and metabolite data demonstrate rapid shifts and subsequent stabilization after an abrupt dietary change in healthy adult dogs. *Anim*. Microbiome 4, 46 (2022). 10.1186/s42523-022-00194-9

6. Lyu, Y. et al. Faecal metabolome responses to an altered dietary protein:carbohydrate ratio in adult dogs. Vet. Q. 43, 1–10 (2023). 10.1080/01652176.2023.2273891

7. Geary, E.L., Oba, P.M., Templeman, J.R. & Swanson, K.S. Apparent total tract nutrient digestibility of frozen raw, freeze-dried raw, fresh, and extruded dog foods and their effects on serum metabolites and fecal characteristics, metabolites, and microbiota of healthy adult dogs. *Transl*. Anim. Sci. 8, txae163 (2024). 10.1093/tas/txae163

8. Hiney, K. et al. Fecal microbiota composition, serum metabolomics, and markers of inflammation in dogs fed a raw meat-based diet compared to those on a kibble diet. Front. Vet. Sci. 11, 1328513 (2024). 10.3389/fvets.2024.1328513

9. Nearing, J.T., et al. Gut microbiome-metabolome interactions during varied low-carbohydrate food consumption. Proc. Natl Acad. Sci. USA 123, e2533462123 (2026). 10.1073/pnas.2533462123

10. Coelho, L.P. et al. Similarity of the dog and human gut microbiomes in gene content and response to diet. Microbiome 6, 72 (2018). 10.1186/s40168-018-0450-3

11. Asaduzzaman, M. et al. Longitudinal long-read microbiome profiling in a canine model reveals how age, diet, and birth mode shape gut community dynamics. mSystems 11, e01279–25 (2026). 10.1128/msystems.01279-25

12. Boldogkői, Z. & Tombácz, D. The canine gut microbiome as a translational model for human health and aging. mBio e01951–26 (2026). 10.1128/mbio.01951-26

13. David, L.A. et al. Diet rapidly and reproducibly alters the human gut microbiome. Nature 505, 559–563 (2014). 10.1038/nature12820

14. Mahieu, N.G. & Patti, G.J. Systems-level annotation of a metabolomics data set reduces 25,000 features to fewer than 1,000 unique metabolites. Anal. Chem. 89, 10397–10406 (2017). 10.1021/acs.analchem.7b02380

15. Sumner, L.W. et al. Proposed minimum reporting standards for chemical analysis: Chemical Analysis Working Group (CAWG), Metabolomics Standards Initiative (MSI). Metabolomics 3, 211– 221 (2007). 10.1007/s11306-007-0082-2

