## Supplementary Figure 1 for "Rapid and largely reversible shifts in the canine fecal metabolome during dietary change"

Expanded pedigree of the nine study dogs

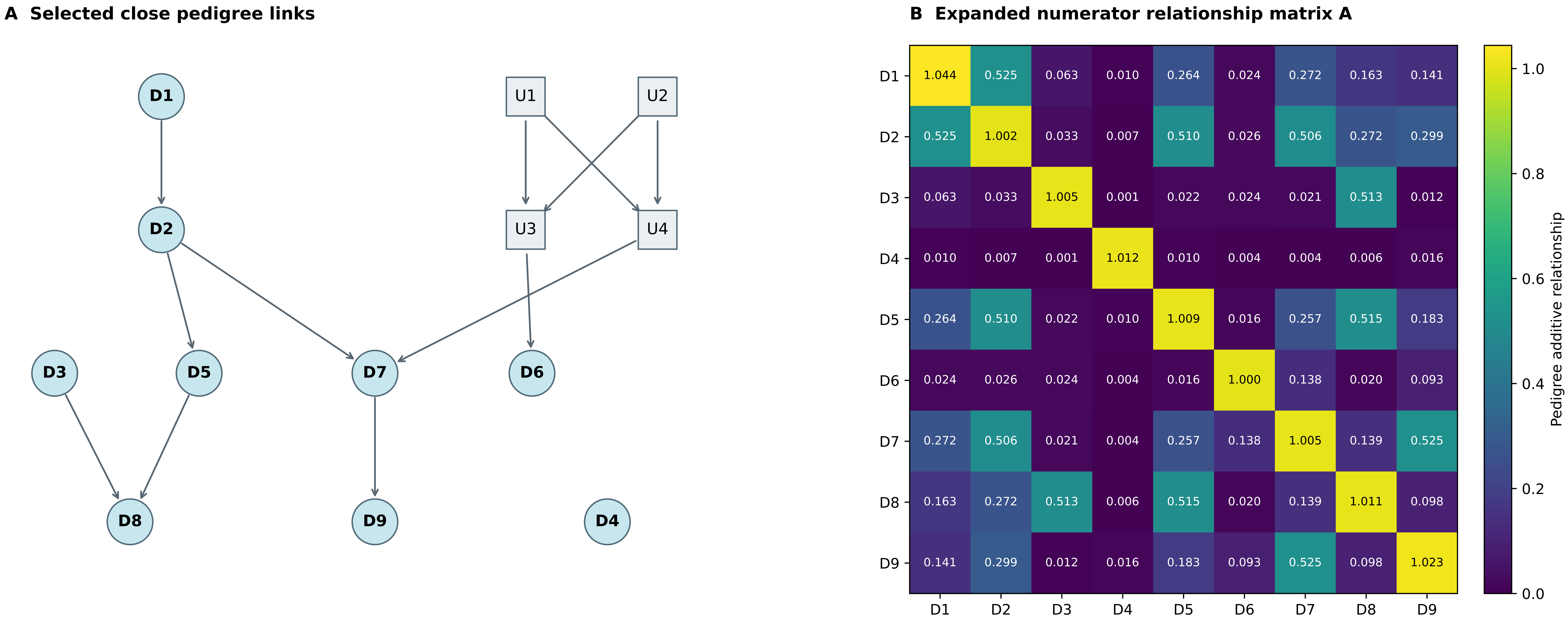


U1 / U2: shared grandparents of D6 and D7
U3: D6 dam; U4: D7 sire (full siblings)
Arrows: parent → offspring

9 study dogs; 238 animals including external ancestors.
9 × 8 / 2 = 36 pairwise comparisons; all Aij > 0.
A diagonal = 1 + F; kinship matrix = A / 2.
