## Supplementary figures and images for "Rapid and largely reversible shifts in the canine fecal metabolome during dietary change"

### Supplementary Figure 2

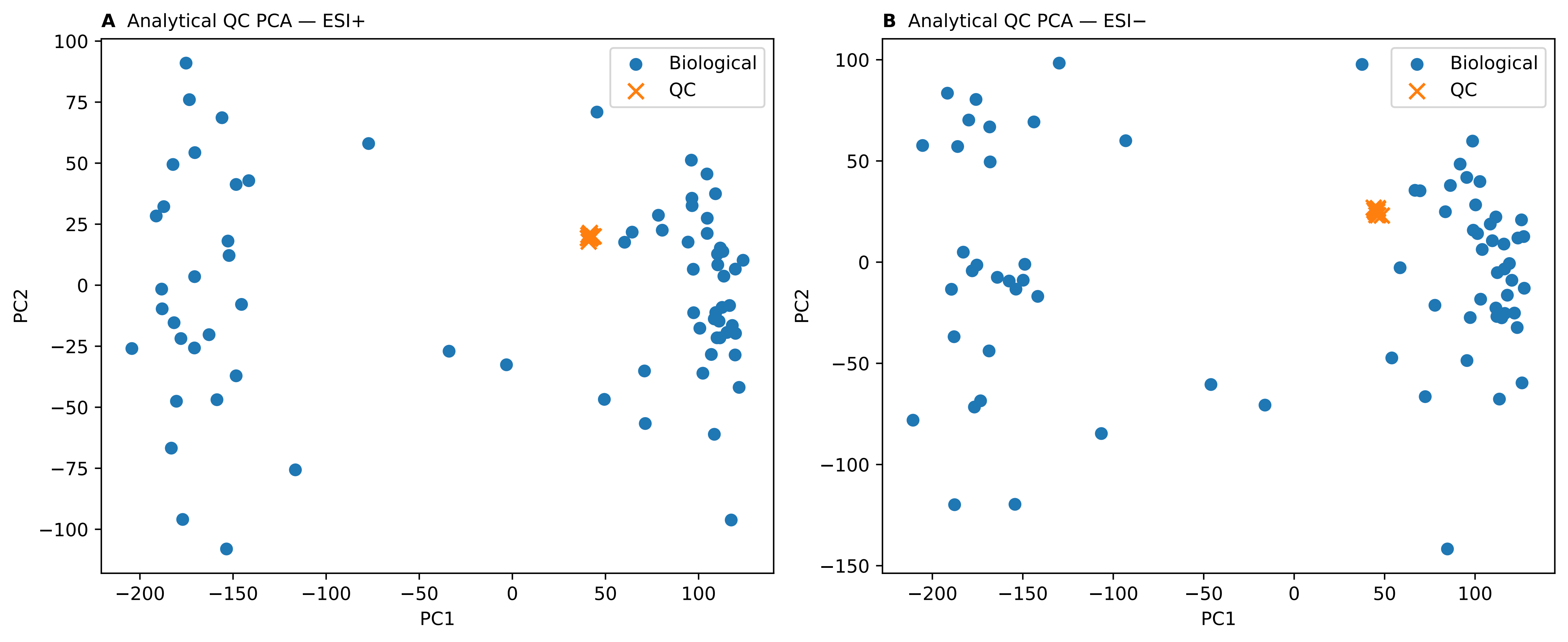

### Supplementary Figure 3

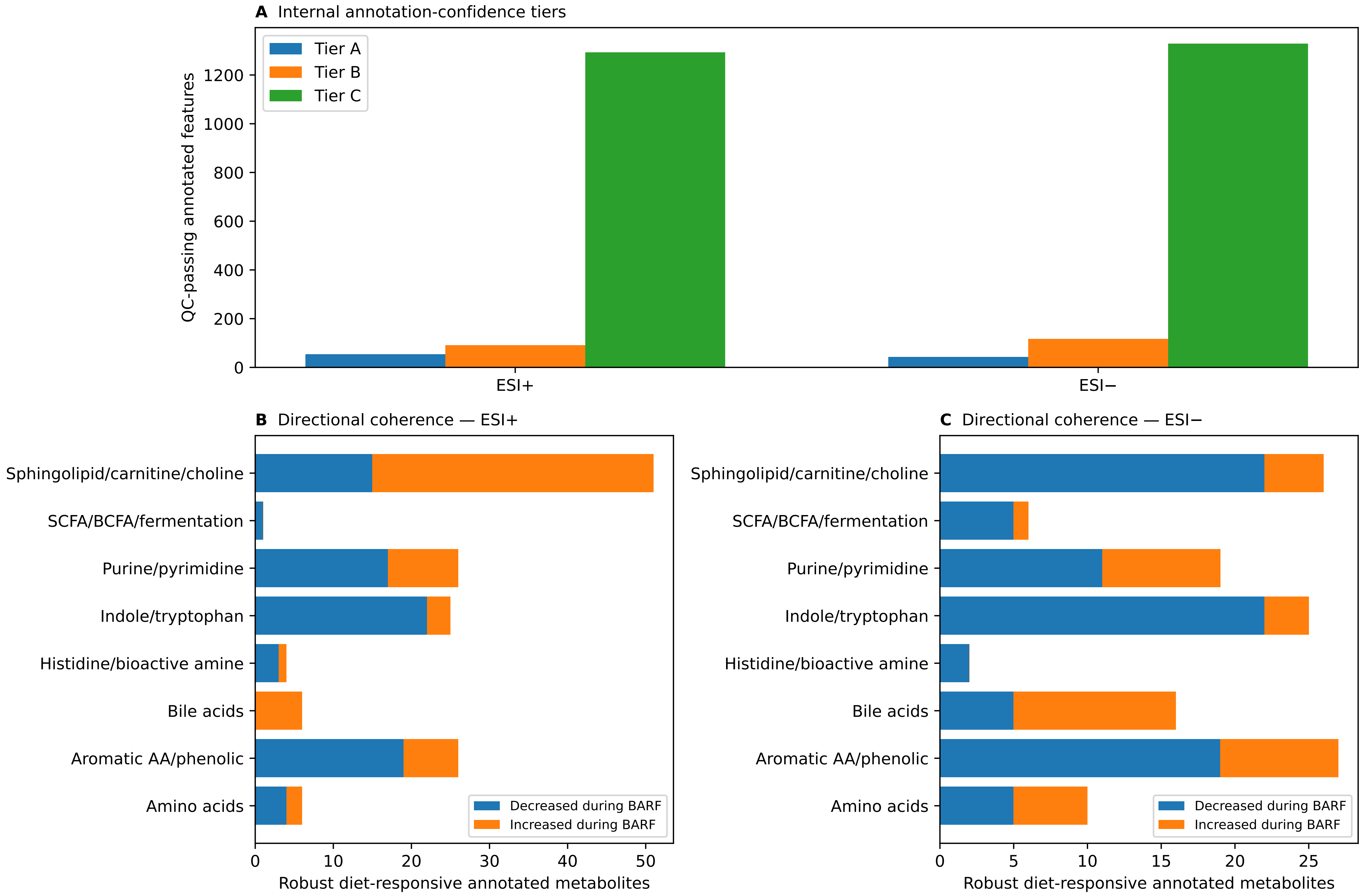

### Supplementary Figure 4

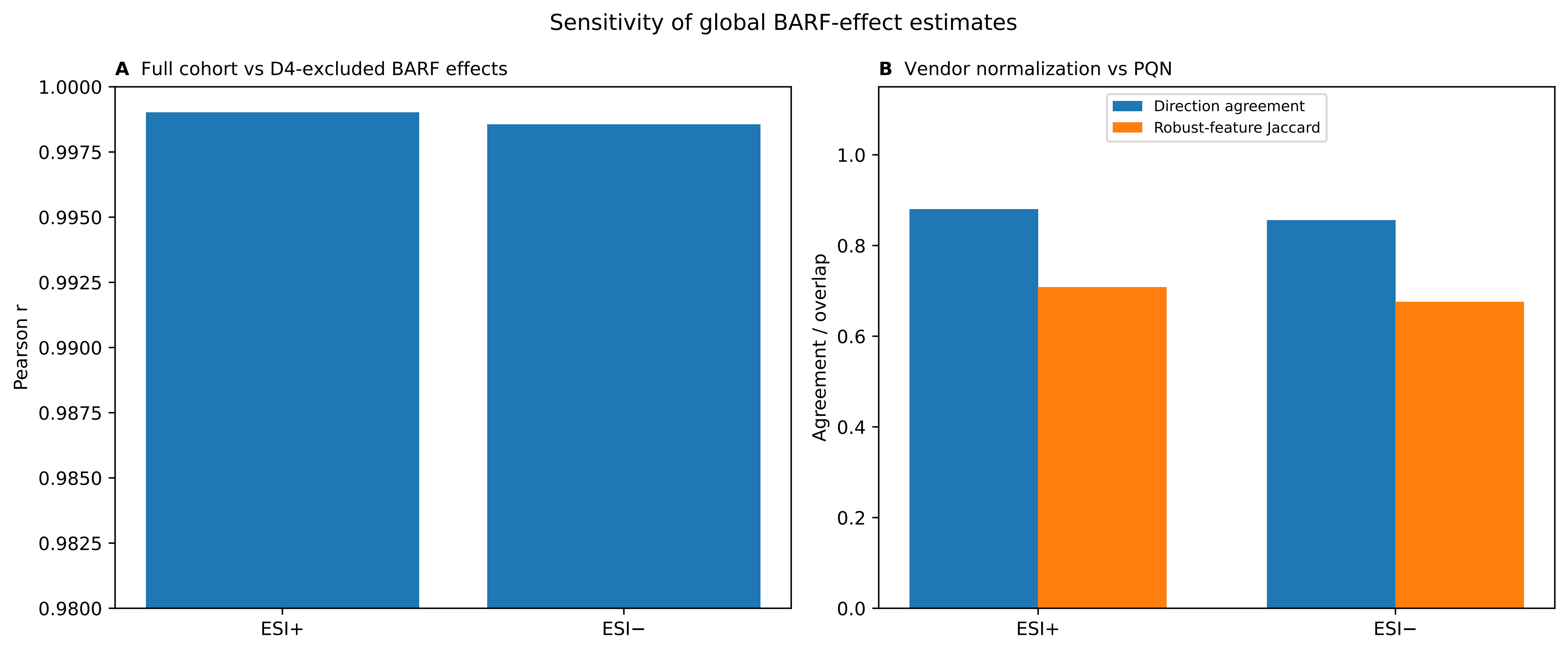

### Supplementary Figure 5

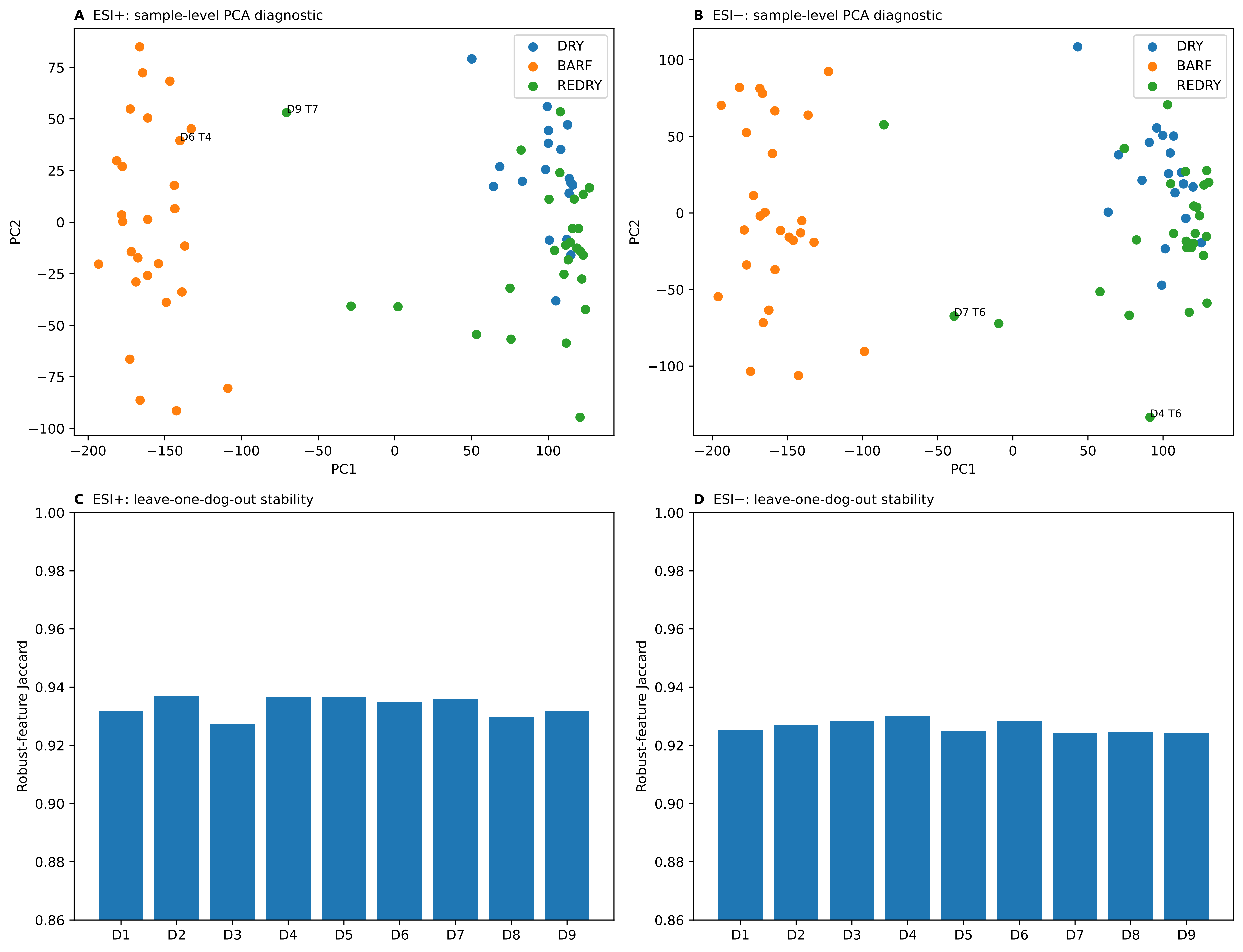

### Supplementary Figure 6

Exploratory D4 peripartum/postpartum trajectories in selected core metabolites

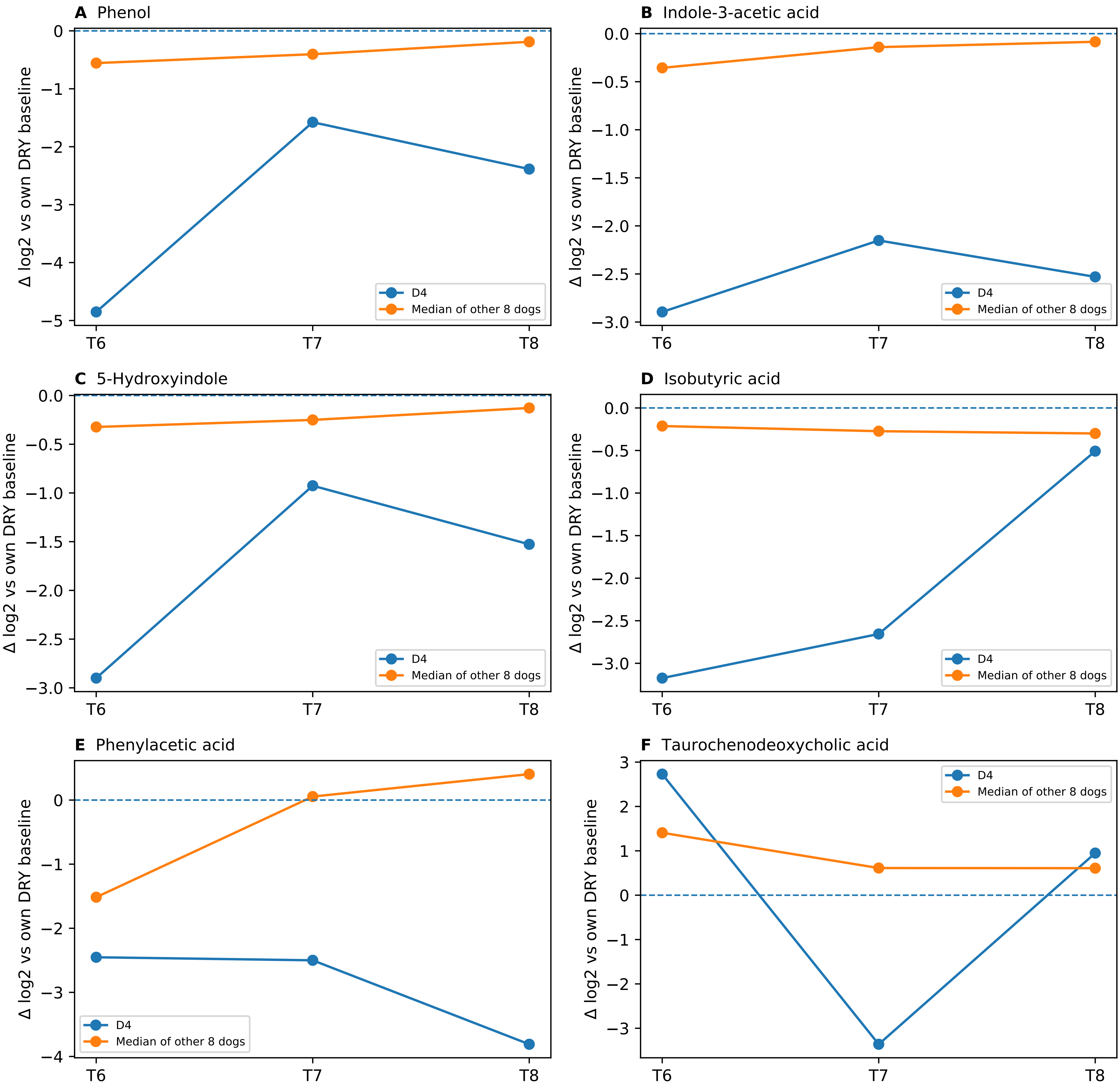

### Supplementary Figure 7

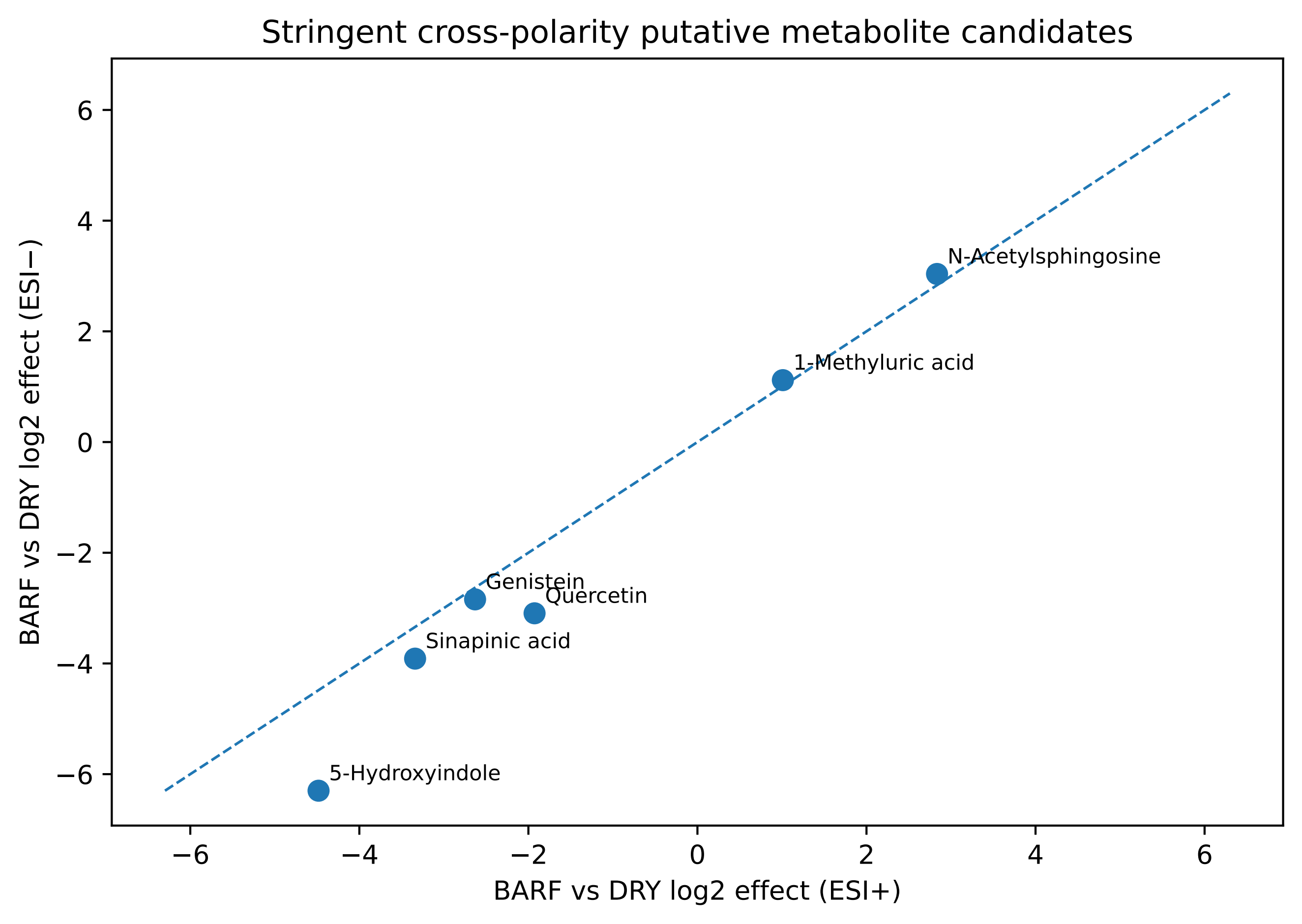

### Supplementary Figure 8

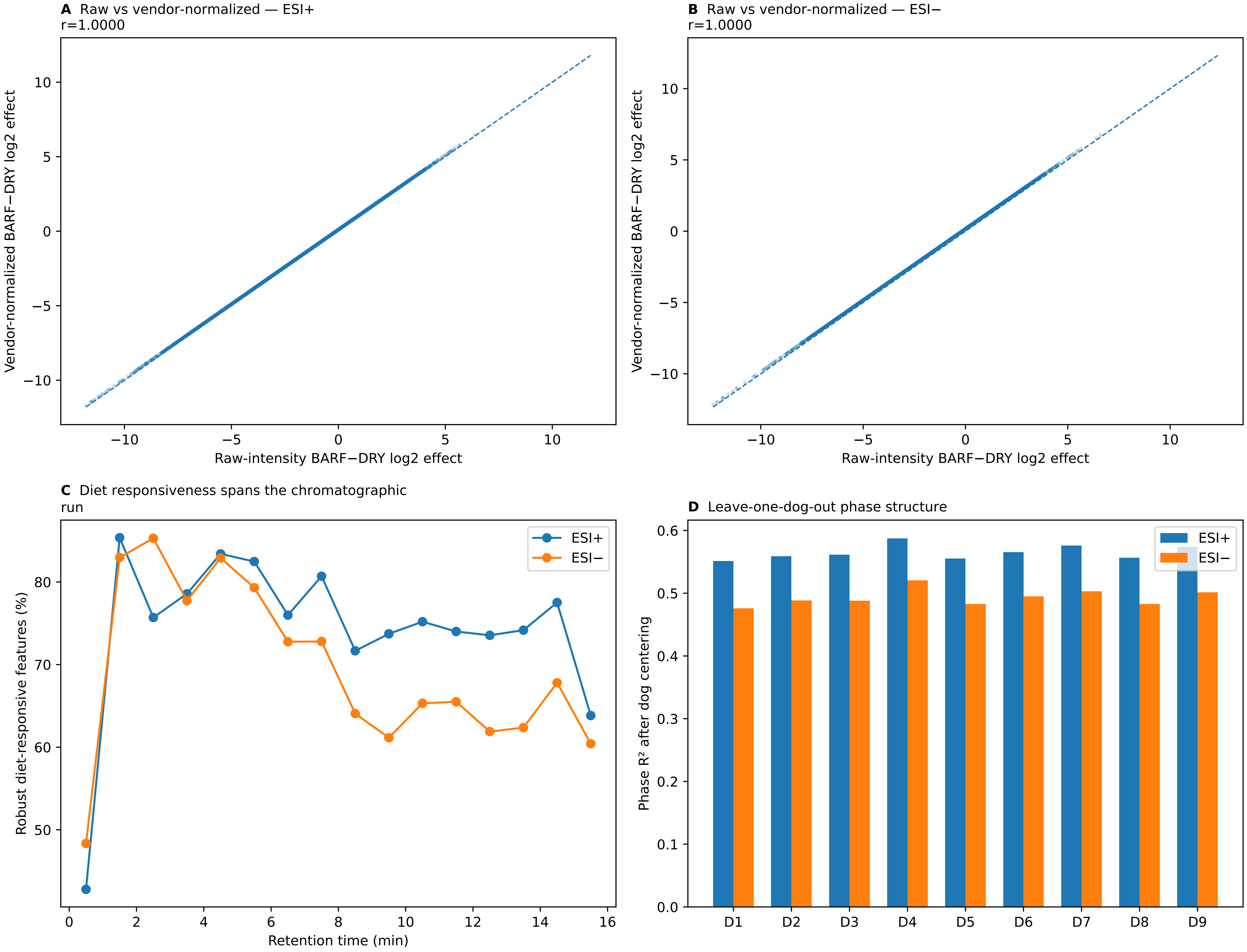

### Supplementary Figure 9

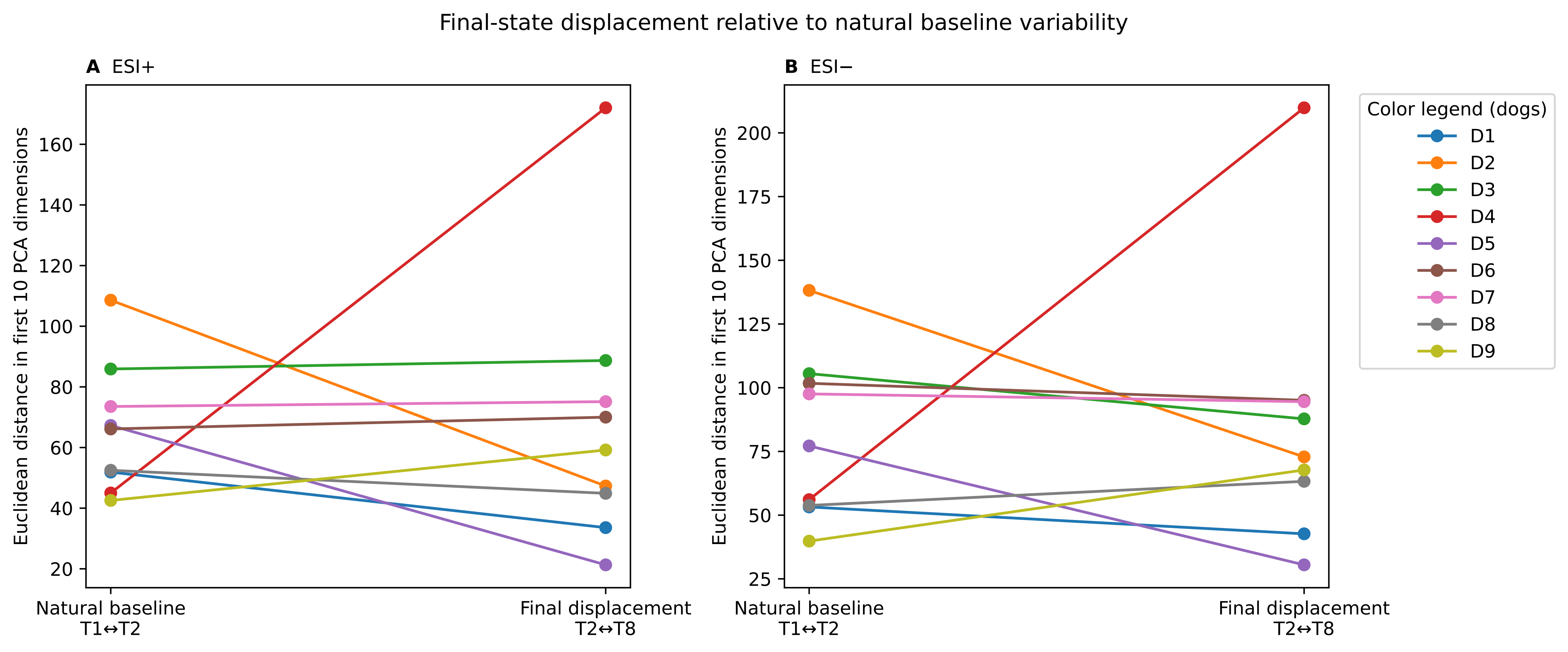

### Supplementary Figure 10

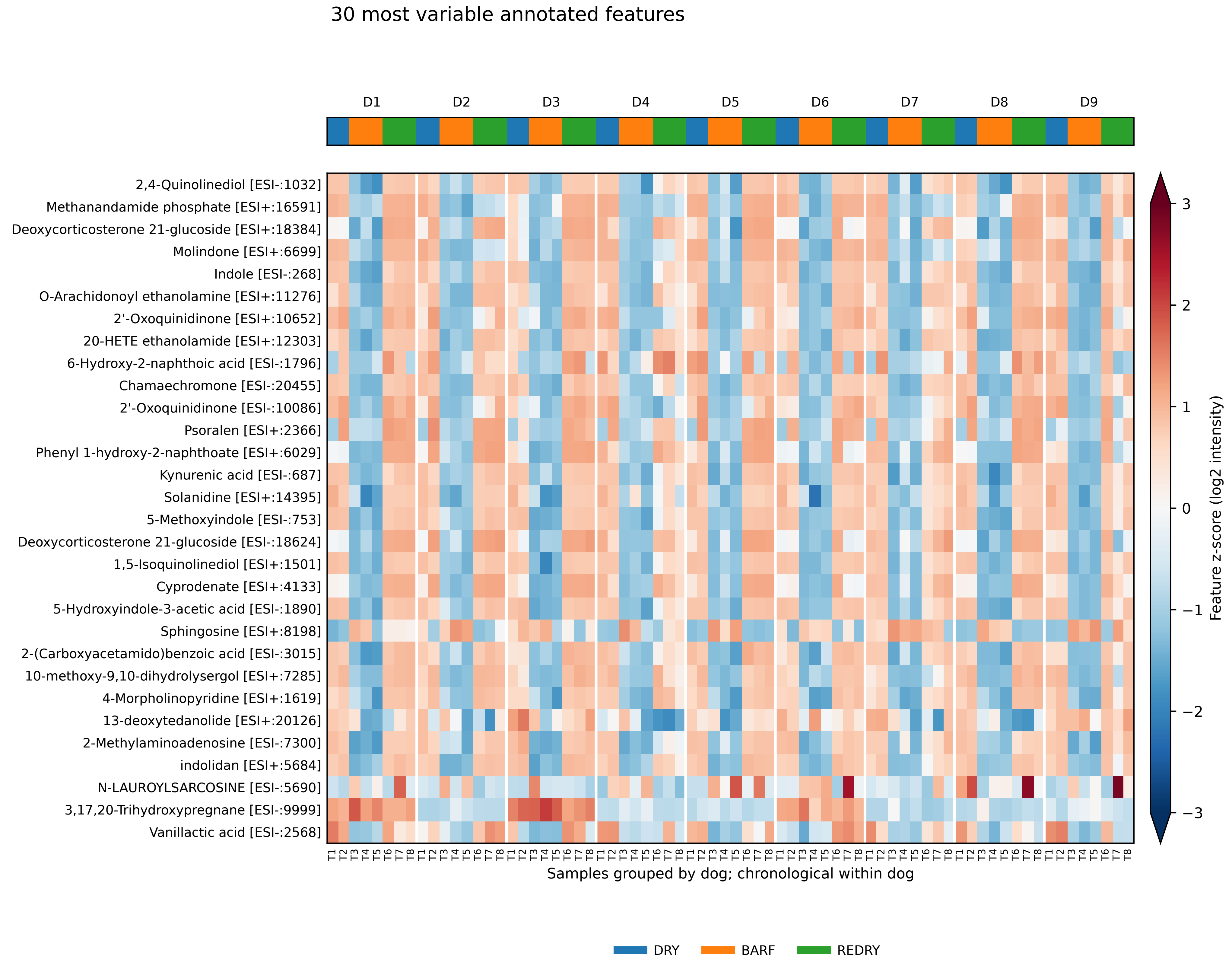

### Supplementary Figure 11

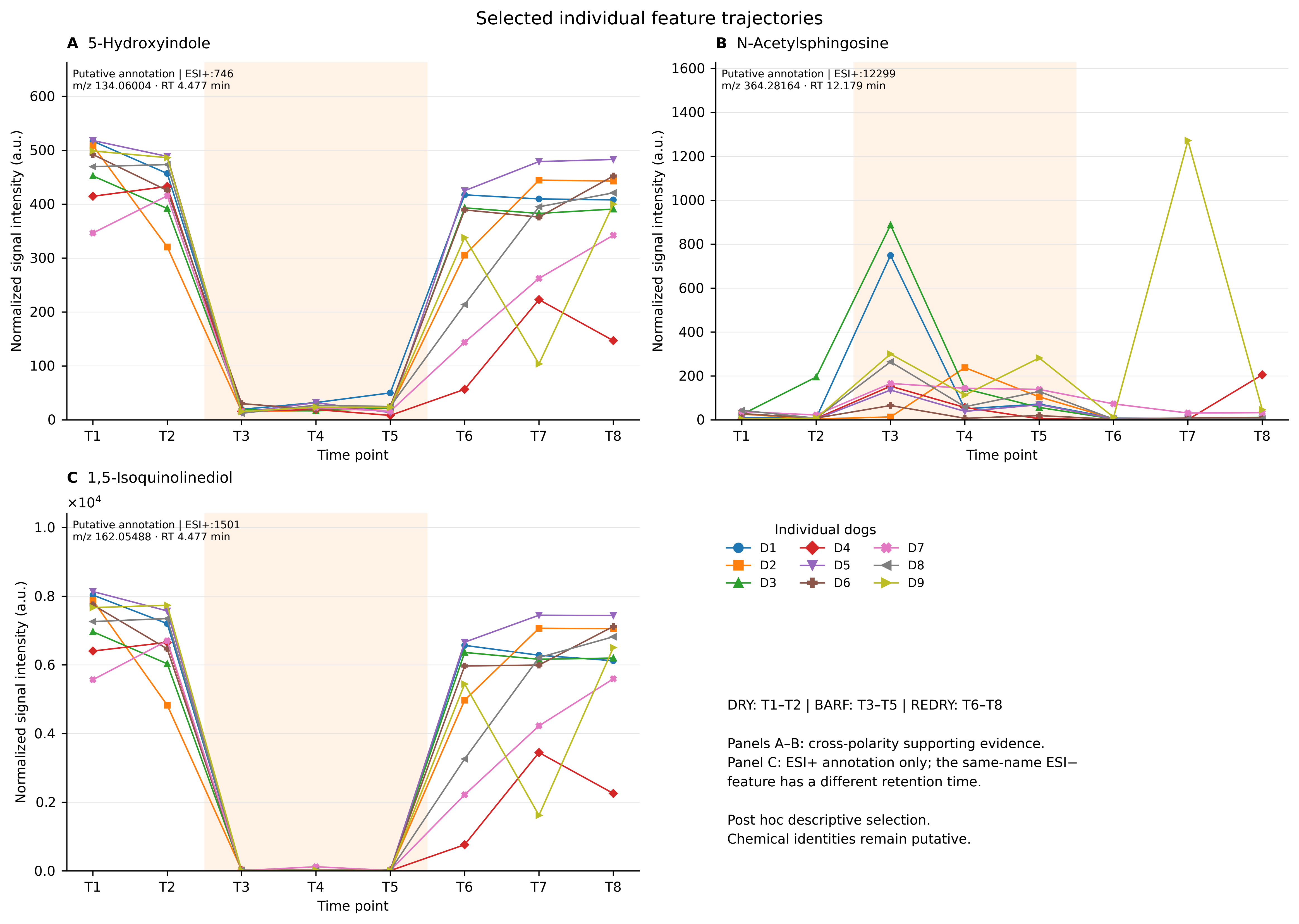
