## Supplementary Legends for "Rapid and largely reversible shifts in the canine fecal metabolome during dietary change"

### Supplementary figure legends

Supplementary Figure S1 | Expanded pedigree and additive relationships. (A) Selected close parent–offspring links among study dogs D1–D9 and four external ancestors. U1 and U2 are Borgarmarken’s Mustang Cobra and Okoska Koroscenko; their full-sibling offspring U3 (Mariell De Marque Lotus Av Fylgia) and U4 (Mariell De Marque Loremo Av Fylgia) are the dam of D6 and sire of D7, respectively. Arrows indicate parentage. Other external parents and distant paths are omitted from this schematic. (B) The study-dog numerator relationship matrix A calculated from the complete 238-animal merged transcription. The diagonal equals 1 + F and kinship equals A/2. The nine study dogs yield 36 distinct pairwise comparisons (9 × 8 / 2), all with documented shared ancestry. Unknown-parent founders were assumed unrelated and noninbred. Tables S15a–c provide the full parentage transcription, matrix, and identifier reconciliation.

Supplementary Figure S2 | Analytical QC in PCA space. (A,B) Pooled QC injections relative to biological samples in ESI+ and ESI−, respectively. QC injections cluster tightly relative to the biological samples. QC intensities were normalized to the full-matrix total signal before applying the feature QC filter.

Supplementary Figure S3 | Annotation confidence and biochemical directionality. (A) Numbers of QC-passing named annotations assigned to internal confidence Tiers A–C in each polarity. (B,C) Directional coherence of selected biochemical classes among robust diet-responsive annotated metabolites in ESI+ and ESI−, respectively.

Supplementary Figure S4 | Sensitivity of BARF-effect estimates. (A) Correlation between full-cohort and D4-excluded effects. (B) Direction agreement and robust-feature Jaccard overlap between vendor normalization and PQN under the harmonized exact sign-flip screen. Near-perfect rank correlation does not imply agreement of absolute effects or selected-feature sets.

Supplementary Figure S5 | Sample-level diagnostics and leave-one-dog-out stability. (A,B) Sample-level PCA diagnostics for ESI+ and ESI−, with flagged phase-specific outliers labeled by study ID and time point. (C,D) Robust-feature Jaccard overlap after leave-one-dog-out analysis in ESI+ and ESI−, respectively.

Supplementary Figure S6 | Exploratory D4 peripartum/postpartum trajectories. D4 trajectories are compared with the median of the other eight dogs for (A) phenol, (B) indole-3-acetic acid, (C) 5-hydroxyindole, (D) isobutyric acid, (E) phenylacetic acid, and (F) taurochenodeoxycholic acid across T6–T8.

Supplementary Figure S7 | Stringent cross-polarity putative metabolite candidates. Candidates required the same name and formula in ESI+ and ESI−, retention-time difference ≤0.05 min, mzCloud score ≥80 in both modes, robust diet response in both modes, and concordant BARF-effect direction. Six candidates met all criteria: 5-hydroxyindole, genistein, 1-methyluric acid, sinapinic acid, quercetin, and N-acetylsphingosine. These remain putative annotations rather than confirmed chemical identities.

Supplementary Figure S8 | Technical robustness analyses. (A,B) Concordance between raw-intensity and vendor-normalized BARF–DRY effect sizes in ESI+ and ESI−. (C) Distribution of robust diet-responsive features across retention time. (D) Leave-one-dog-out stability of phase-associated multivariate structure after dog centering. Phase-associated variance was calculated in the log2-transformed, Pareto-scaled full-feature space; scaling was estimated in the full cohort.

Supplementary Figure S9 | Final-state displacement relative to natural baseline variability. (A,B) For each dog, T2→T8 distance was compared with T1→T2 baseline distance in the first 10 PCA dimensions for ESI+ and ESI−, respectively. Line colors identify D1–D9 as shown in the color legend; exact paired P=0.906 in ESI+ and P=0.883 in ESI−. This comparison does not test equivalence with baseline.

Supplementary Figure S10 | Individual-sample patterns among the 30 most variable annotated LC–MS features. Features were ranked across both polarities by the variance of log2 vendor-normalized intensity over 72 samples, after intersection of the annotated master table with the raw-QC RSD <30% feature set. Rows denote measured features, identified by polarity and vendor feature number; chemical names are putative annotations and repeated names do not establish distinct metabolites. Columns are grouped by study dog D1–D9 and ordered T1–T8 within each dog. The upper strip marks DRY (T1–T2), BARF (T3–T5) and REDRY (T6–T8). Colors show feature-wise z-scores calculated over all 72 log2 intensities using the sample standard deviation; values beyond ±3 are displayed at the color limits without changing the source values. Selection and display use the same cohort and are descriptive rather than independent validation. Full feature names and selection metadata are supplied in Table S15.

Supplementary Figure S11 | Selected individual feature trajectories. Panels show vendor-normalized ESI+ signal intensities for (A) feature 746, putatively annotated as 5-hydroxyindole; (B) feature 12299, putatively annotated as N-acetylsphingosine; and (C) feature 1501, putatively annotated as 1,5-isoquinolinediol. Each line denotes one dog (D1–D9); the shaded region marks BARF sampling points T3–T5. The T1–T8 axis denotes successive sampling occasions rather than equal elapsed-time intervals. Panels A and B were selected from the stringent cross-polarity candidate set (Table S13) to illustrate opposite directions of response. Panel C retains a previously explored trajectory with an mzCloud score of 89.8 in ESI+; its same-name ESI− annotation has a different retention time and was not treated as corroboration of chemical identity. This post hoc descriptive selection does not provide independent confirmation of the response or chemical identity. Intensities are arbitrary units, not absolute concentrations; all plotted values are positive, without additional imputation or zero replacement. Feature metadata are supplied in Table S16.

### Supplementary table legends

Table S1a | Anonymized dog metadata. Age, sex, housing, and study identifiers D1–D9.

Table S1b | Anonymized sample metadata. Sampling time point, diet phase, and D4 reproductive-status notes.

Table S2 | QC-filtered annotated-metabolite master table. Annotation-confidence tier, within-dog BARF effect, FDR, recovery class, and cross-polarity corroboration for named QC-passing features. The inclusion filter is based on vendor-normalized QC intensities, whereas global feature analyses use raw-QC RSD; four rows pass the former threshold but not the latter.

Table S3 | Cross-polarity matches. Putative metabolites matched across ESI+ and ESI− by name, formula, and retention time.

Table S4 | Exploratory biochemical-class directionality. Increased and decreased robust annotated feature counts, two-sided exact binomial p values, and BH-adjusted q values across eight classes per polarity. Correlation and redundancy among LC–MS signals are not modeled.

Table S5a | Follow-up mixed-effects models for all 13 candidate metabolites, including N-acetyltryptophan. BARF-versus-DRY and REDRY-versus-DRY effects, BH-adjusted q values across the 13-candidate family, fitting diagnostics, and D4-exclusion estimates. Reproduced standard errors and 95% asymptotic Wald confidence intervals are appended for the full-cohort models. The main display panel contains the 12 candidates meeting the BARF criterion; models are not independent validation.

Table S5b | Expanded-pedigree sensitivity models for all 13 candidate metabolites. The numerator relationship matrix was rebuilt from the merged 238-animal pedigree, and pedigree, independent-dog and residual variance components were re-estimated by profile maximum likelihood. The table includes effects, standard errors, asymptotic two-sided Wald p values and 95% confidence intervals, BH-adjusted q values across 13 candidates per contrast, convergence and variance-boundary diagnostics. A boundary variance estimate is not evidence of absent biological genetic variation.

Table S6a | Normalization sensitivity under a harmonized exact sign-flip screen. Rank correlation, direction agreement, robust-feature counts and Jaccard overlap, and the log2 effect shift are reported. PQN divides raw QC-passing intensities by sample-wise median quotients relative to the featurewise median reference across all 72 biological samples.

Table S6b | Global BARF-effect sensitivity after exclusion of D4, summarized separately by polarity. Candidate-specific D4-exclusion results are supplied in Table S5a.

Table S6c | Influence of flagged sample removal on the BARF-versus-DRY contrast recomputed from within-dog phase means. Removal of a REDRY-only sample leaves this contrast unchanged.

Table S6d | Leave-one-dog-out sensitivity using the harmonized exact sign-flip screen. Eight-dog analyses enumerate 256 sign configurations and require concordant direction in at least seven dogs. Jaccard overlap compares each eight-dog feature set with the full nine-dog set.

Table S7 | D4 peripartum/postpartum exploratory results. Core-metabolite trajectories and D4-specific departures during T6–T8.

Table S8 | Ion/adduct mass-offset sanity checks. Consistency of selected core putative annotations with plausible common ion/adduct forms.

Table S9 | Extended transition summary. Significant feature counts and effect summaries for T1→T2, T2→T3, T5→T6, and T2→T8; separate fields report the shared significant feature count, opposite-direction percentage, and cosine similarity. Recovery summaries are included in the overlap rows; blank fields denote nonapplicable metrics.

Table S10 | Temporal-cluster summary. Cluster sizes and mean centroids for the three dominant response trajectories in each polarity.

Table S11 | Raw-normalization reconstruction and sensitivity. Exact reconstruction of vendor total-signal normalization and concordance of raw and normalized BARF effects.

Table S12 | Final-state versus natural-baseline multivariate distances. Dog-level T1→T2 and T2→T8 distances in the first 10 PCA dimensions.

Table S13 | Stringent cross-polarity candidate list. Putative metabolite candidates satisfying cross-polarity name, formula, retention-time, mzCloud, robust-response, and direction-concordance criteria.

Table S14a | Expanded parentage transcription. All 238 unique animals represented in the merged pedigree, with sire, dam, source screenshot, inferred sex where available, founder assumption and pedigree inbreeding coefficient F. Blank parent fields indicate missing information, not proven unrelatedness. Registered-name spelling and identity links remain subject to author verification against original registry records.

Table S14b | Expanded numerator relationship matrix A for study dogs D1–D9. Coefficients use all documented parentage paths in Table S14a; Aii = 1 + Fi, and the kinship matrix equals A/2. Values depend on the assumption that unknown-parent founders are unrelated and noninbred.

Table S14c | Study-dog identifier reconciliation and selected shared-ancestor registration identifiers. Dataset names, manuscript names, registered names and screenshot evidence are provided; Cuidado Xotic supersedes the inconsistent Quisite label following the author’s correction.

Table S15 | Selection metadata for the Top30 individual-sample heatmap. Exactly 30 mode-specific feature identifiers ranked by log2 variance, with putative annotation, retention time, mass-to-charge ratio, existing annotated-table statistics and selection rank. The QC_RSD_percent field is inherited from Table S2 (normalized QC); each selected row also passes the raw-QC filter. The table is a descriptive selection record, not a new set of hypothesis tests.

Table S16 | Feature metadata for selected individual trajectories. Panel, polarity, vendor feature number, putative name, m/z, retention time, annotation scores and existing annotated-table statistics for Supplementary Figure S11. Selection is post hoc and descriptive; no new hypothesis tests are introduced.
